# TIGAR-V2: Efficient TWAS Tool with Nonparametric Bayesian eQTL Weights of 49 Tissue Types from GTEx V8

**DOI:** 10.1101/2021.07.16.452700

**Authors:** Randy L. Parrish, Greg C. Gibson, Michael P. Epstein, Jingjing Yang

**Affiliations:** Center for Computational and Quantitative Genetics, Department of Human Genetics, Emory University School of Medicine, Atlanta, GA, 30322, USA; School of Biology, Georgia Institute of Technology, Atlanta, GA, 30332, USA

**Keywords:** Transcriptome-wide association study, TWAS, GWAS, cis-eQTL, Nonparametric Bayesian DPR, Elastic-Net, TIGAR

## Abstract

Standard Transcriptome-Wide Association Study (TWAS) methods first train gene expression prediction models using reference transcriptomic data, and then test the association between the predicted genetically regulated gene expression and phenotype of interest. Most existing TWAS tools require cumbersome preparation of genotype input files and extra coding to enable parallel computation. To improve the efficiency of TWAS tools, we develop TIGAR-V2, which directly reads VCF files, enables parallel computation, and reduces up to 90% computation cost (mainly due to loading genotype data) compared to the original version. TIGAR-V2 can train gene expression imputation models using either nonparametric Bayesian Dirichlet Process Regression (DPR) or Elastic-Net (as used by PrediXcan), perform TWAS using either individual-level or summary-level GWAS data, and implements both burden and variance-component statistics for gene-based association tests. We trained gene expression prediction models by DPR for 49 tissues using GTEx V8 by TIGAR-V2 and illustrated the usefulness of these Bayesian cis-eQTL weights through TWAS of breast and ovarian cancer utilizing public GWAS summary statistics. We identified 88 and 37 risk genes respectively for breast and ovarian cancer, most of which are either known or near previously identified GWAS (~95%) or TWAS (~40%) risk genes and three novel independent TWAS risk genes with known functions in carcinogenesis. These findings suggest that TWAS can provide biological insight into the transcriptional regulation of complex diseases. TIGAR-V2 tool, trained Bayesian cis-eQTL weights, and LD information from GTEx V8 are publicly available, providing a useful resource for mapping risk genes of complex diseases.

## Introduction

A transcriptome-wide association study (TWAS) ^1–5^ is a popular technique widely used for integrating reference transcriptomic data with GWAS data to conduct gene-based association studies. TWAS has been shown to improve the power of identifying GWAS risk loci as well as illustrate the underlying biological mechanism of GWAS loci, for example in studies of schizophrenia [MIM: 181500] ^6^, age-related macular degeneration [MIM: 603075] ^7^, and broad types of complex traits ^8^. Especially, the risk genes identified by TWAS have genetic effects potentially mediated through gene expression.

The standard two-stage TWAS methods ^1–3^ first fit gene expression prediction models using the reference transcriptomic and genetic data profiled for the same samples, and then test the association between the predicted genetically regulated gene expression (GReX) and phenotype of interest for the test GWAS cohort. The TWAS framework enables the advantages of using publicly available reference transcriptomic data such as the Genotype-Tissue Expression (GTEx) project ^9,10^ and summary-level GWAS data ^11,12^.

However, most of the existing tools ^1,2,5^ require cumbersome preparation of genotype data files and fail to take advantage of parallel computing to improve computational efficiency. These result in difficulties for users who need to train gene expression prediction models by using their own reference transcriptomic and genetic data. Here, we develop a new version of the Transcriptome-Integrated Genetic Association Resource (referred to as TIGAR-V2) that takes genotype data of the Variant Call Format (VCF) as input, conducts 5-fold cross-validation ^13^ to evaluate trained gene expression prediction models, and enables parallel computation to take advantage of high performance computing clusters.

Additionally, TIGAR-V2 can train gene expression imputation models using either nonparametric Bayesian Dirichlet Process Regression (DPR) ^14^ or Elastic-Net penalized regression (as used by PrediXcan ^1^). TIGAR-V2 can perform TWAS using either individual-level or summary-level GWAS data. Besides the burden type TWAS test ^1^, the software further implements an additional variance-component test ^15^ for TWAS that retains power under model misspecification.

To make TIGAR-V2 a convenient resource for the public, we trained nonparametric Bayesian DPR gene expression prediction models for 49 tissues from the GTEx V8 reference panel ^10^. These estimated tissue-specific SNP effect sizes on the expression quantitative traits (eQT) are considered as Bayesian eQTL weights per gene and are provided along with this TIGAR-V2 tool, which can be conveniently used for follow-up gene-based association studies using both individual-level and summary-level GWAS data. In our example application studies, we used eQTL weights obtained from transcriptomic data of breast mammary tissue and ovary tissue from the GTEx V8 reference panel along with publicly available GWAS summary statistics ^11,12^ to conduct TWAS for studying breast cancer [MIM: 114480] and ovarian cancer [MIM: 167000].

In the following sections, we first outline the TIGAR-V2 framework. We then describe the application of TIGAR-V2 to train gene expression prediction models with the GTEx V8 reference data and TWAS of breast cancer and ovarian cancer. Model training and application results are described. Finally, we conclude with a discussion.

## Material and Methods

### TIGAR-V2 framework

#### Gene expression prediction model

The standard two-stage TWAS ^1–3^ first fits gene expression prediction models by taking genotype data (**G**) of cis-SNPs (within ±1MB of the target gene *g* ^2,16,17^) as predictors, assuming the following additive genetic model for the expression quantitative trait (**E**_*g*_) with respect to a target gene *g*,

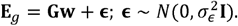

The cis-eQTL effect size vector **w** can be estimated by different regression methods from the reference (i.e., training) data. For example, PrediXcan estimates **w** by a general linear regression model with Elastic-Net penalty ^1^; FUSION estimates **w** by Elastic-Net, LASSO^18^, linear mixed modeling, Sum of Single Effects (SuSiE)^19^, and Bayesian Sparse Linear Mixed Model (BSLMM)^20^; and TIGAR estimates **w** by a nonparametric Bayesian DPR model (Text S1) ^3,14^.

TIGAR-V2 implements both nonparametric Bayesian DPR ^14^ and general linear regression with Elastic-Net penalty as used by PrediXcan ^1^ to estimate **w**, which are eQTL effect sizes in a broad sense not considering whether the SNP has a genome-wide significant eQTL p-value. Additionally, TIGAR-V2 runs 5-fold cross validation ^13^ with the reference data by default to provide an average prediction *R*^2^ per gene across 5 folds of validation data (referred to as 5-fold CV *R*^2^). The 5-fold CV *R*^2^ can be used to evaluate if the trained gene expression prediction model is “valid” for follow-up TWAS (e.g., using the threshold of 5-fold CV *R*^2^ > 0.005). Here, we use a more liberal threshold than the threshold 0.01 used by previous studies ^2,21,22^ to allow more genes to be tested in follow-up TWAS. Because the follow-up gene-based association Z-score test statistic is essentially a weighted average of single variant GWAS Z-score statistics with variant weights provided by the eQTL effect sizes (Equations 2 and 3), poorly estimated eQTL weights would only reduce power but will not increase false positive rate under null hypothesis.

#### Gene-based association study

With the estimates of cis-eQTL effect sizes **ŵ** and individual-level GWAS data of test samples, TIGAR-V2 predicts GReX values by taking estimates of cis-eQTL effect sizes (outputs from the step of training gene expression prediction models) and genotype data (VCF files, G_test_) of test samples as inputs, and using formula 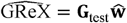. TIGAR-V2 implements the burden^23,24^ type TWAS test by testing the association between 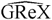 and the phenotype of interest (PED format) based on the general linear regression model, with the phenotype as response variable and predicted 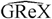 as a test covariate (Text S2.1). TIGAR-V2 implements the variance-component TWAS test by ^15^ using the sequence kernel association test (SKAT) framework ^25^ with variant weights provided by eQTL effect size estimates **ŵ**. Variance-component TWAS test is recommended if the assumption of the linear relationship between the SNP effect sizes on phenotype and eQTL weights is violated (see Text S2.2). Note that here the eQTL weights **ŵ** are specific to the test gene and specific to the tissue type of the reference transcriptomic data.

With summary-level GWAS data (i.e., Z-score statistic values from single variant GWAS tests) of test samples, TIGAR-V2 tests the gene-based association by using both burden ^23,24^ and variance-component ^15^ test statistics, where cis-eQTL effect size estimates **ŵ** are taken as variant weights.

In particular, for burden test, we found that the FUSION Z-score statistic ^2^ as given by Equation (2) will lead to inflated false positive findings if **ŵ** are estimated using non-standardized reference data (i.e., centered gene expression and genotype data as described in Text S1 for Bayesian DPR model); the S-PrediXcan test statistic ^26^ as given by Equation (3) should be used in this situation instead. We also show that both FUSION and S-PrediXcan test statistics are equivalent if **ŵ** are estimated using standardized reference data (Text S2.2). The S-PrediXcan test statistic is the default test statistic implemented by TIGAR-V2.

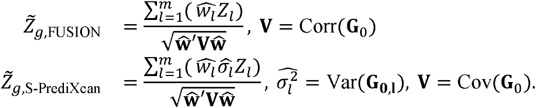

Here, *Z*_*l*_ the Z-score statistic values from single variant GWAS tests (i.e., summary-level GWAS data). The required linkage disequilibrium (LD) covariance matrix (or correlation matrix for FUSION test statistic) among test cis-SNPs (**V**), and the variance of test cis-SNPs 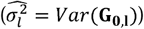 can be obtained from reference genotype data (**G**_0_) such as 1000 genome ^27^ and GTEx V8 ^10^.

#### Tool framework

The tool framework of TIGAR-V2 is shown in Figure 1, where all TWAS steps in TIGAR-V2 are enabled using Python and Bash scripts. Python libraries “pandas” ^28,29^, “numpy” ^28,30^, “scipy” ^31^, “sklearn” ^32,33^, and “statsmodels” ^34^ are used to develop TIGAR-V2. Genotype data in VCF saved as one file per chromosome are input genotype files for TIGAR-V2. TABIX tool ^35^ is used to extract genotype data per target gene efficiently from VCF genotype files. Parallel computation is enabled by using the “multiprocessing” Python library, allowing users to train gene expression prediction models and test gene-based association of multiple genes in parallel.

**Figure 1.**
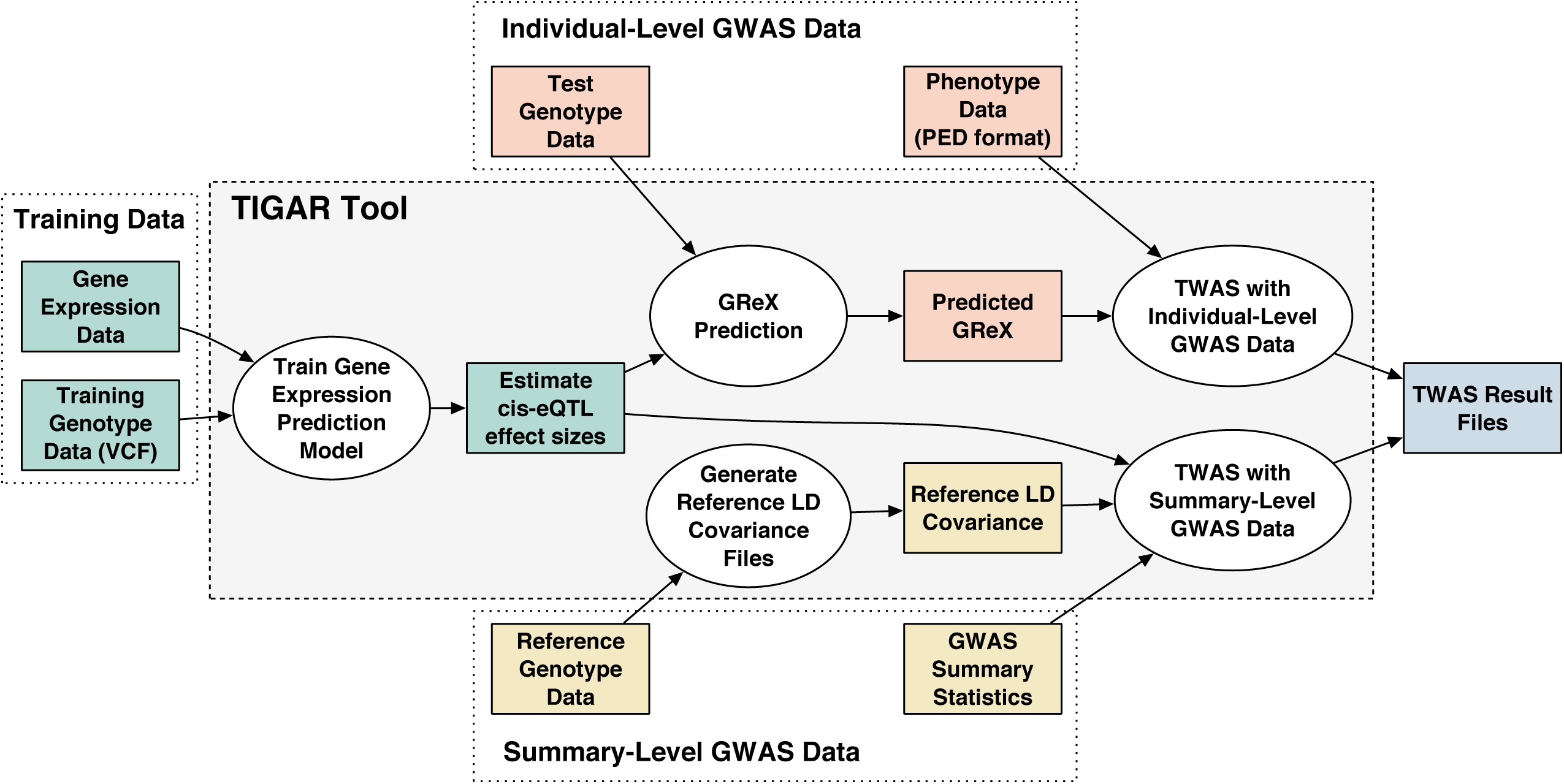
TIGAR-V2 framework. Including TWAS steps of training gene expression prediction models from reference data, predicting GReX with individual-level GWAS data, and testing gene-based association with both individual-level and summary-level GWAS data.

This new version uses fewer Python library dependencies for easier set up, speeds up computation by improving genotype data loading and using functions from the “numpy” Python library, reduces required memory usage by loading genotype data from VCF files with a row-by-row increment, and adds the function to conduct the recently published variance-component gene-based association test ^15^. For example, for training gene expression prediction models by Bayesian DPR method with 129 samples, ~ 1800 SNPs per gene, four genes, and a single core, the computation time is reduced up to 90% and memory usage up to 50% (mainly due to improved genotype data loading from VCF files), compared to the initial TIGAR tool. The memory usage is linear with respect to the number of cis-SNP predictors of the target gene and the training sample size. Training gene expression prediction models using the GTEx V8 reference data requires less than 8GB of memory per gene, with number of test SNPs up to ~10K per gene and training sample size up to ~600. We would suggest users to run one gene per typical computation core in a high-performance computing cluster, e.g., running 4 genes in parallel per chromosome if 4 cores are requested.

### Reference resource from GTEx V8

#### Train Bayesian DPR eQTL weights from GTEx V8

The Genotype-Tissue Expression (GTEx) project V8 (dbGaP phs000424.v8.p2) contains comprehensive profiling of whole genome sequencing (WGS) genotype data and RNA-sequencing (RNA-seq) transcriptomic data (15,253 normal samples) across 54 tissue types of 838 donors ^9,10,36,37^. GTEx V8 provides useful reference data for training tissue-specific gene expression prediction models for a diverse of tissue types on human bodies. Both PrediXcan and FUSION tools use GTEx V8 data as the reference data, and provide estimated cis-eQTL weights per gene with respect to 49 tissue types that have >70 samples with profiled WGS genotype and RNA-seq transcriptomic data (Figure 2A) as a public resource for TWAS.

**Figure 2.**
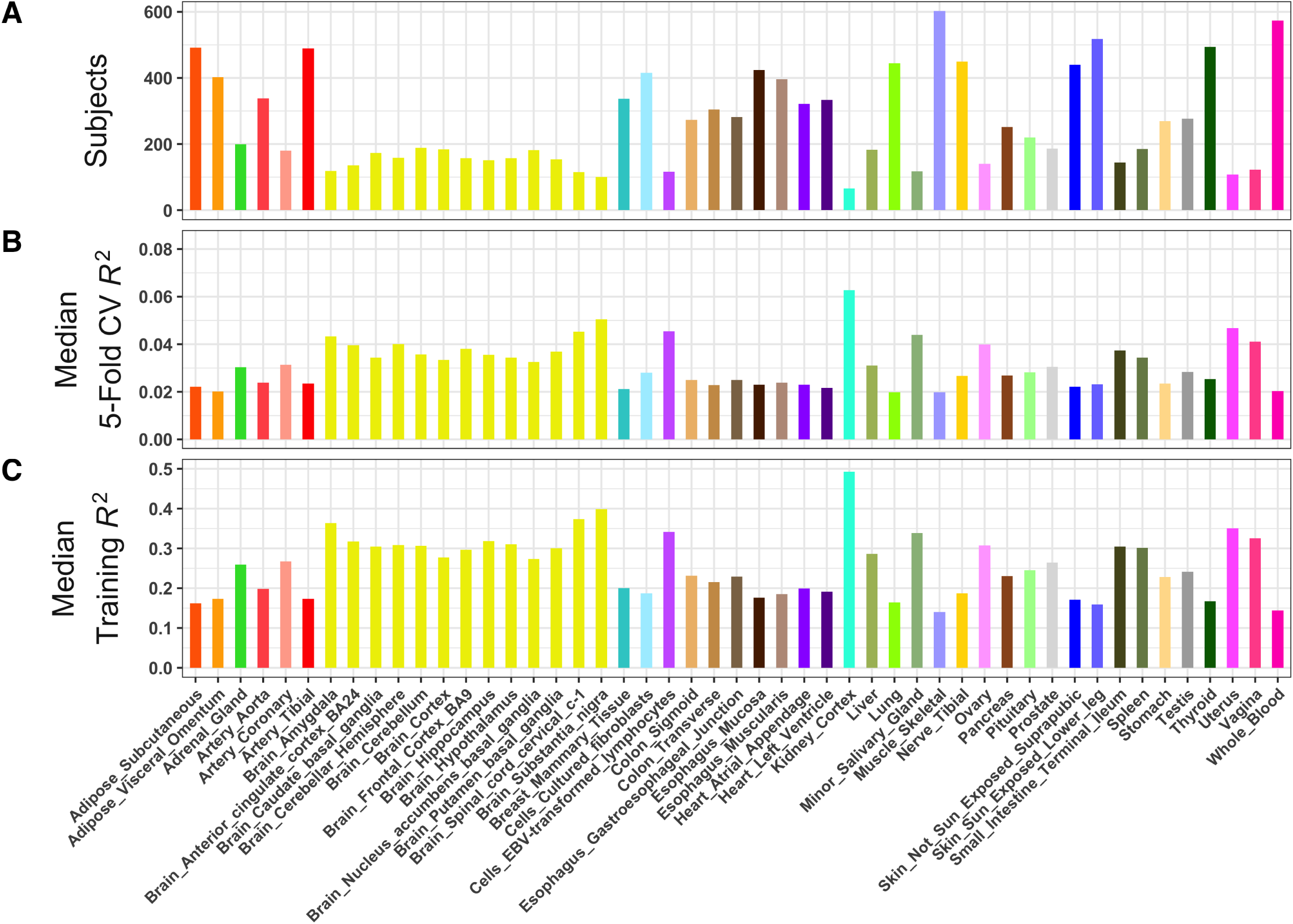
Trained gene expression prediction models of 49 tissue types from GTEx V8 by TIGAR using the nonparametric Bayesian DPR method. (A) Number of training sample size per tissue, (B) Median 5-Fold CV *R*^2^ per tissue, and (C) Median Training *R*^2^ per tissue. Colors are coded with respect to groups of tissue types.

Here, we also train tissue-specific gene expression prediction models for these 49 tissue types using the nonparametric Bayesian DPR method previously implemented in TIGAR. WGS genotype data of cis-SNPs within ±1MB around gene transcription start sites (TSS) of the target gene were used as predictors. In particular, variants with missing rate < 20%, minor allele frequency > 0.01, and Hardy-Weinberg equilibrium p-value > 10^−5^ were considered for fitting the gene expression prediction models. Gene expression data of Transcripts Per Million (TPM) per sample per tissue were downloaded from the GTEx portal. Genes with > 0.1 TPM in ≥ 10 samples were considered. Raw gene expression data (TPM) were then adjusted for age, body mass index (BMI), top five genotype principal components, and top probabilistic estimation of expression residuals (PEER) factors ^38^. The gene expression data of breast mammary tissue were further adjusted for *ESR1* expression following previous TWAS analysis of breast cancer ^39^.

Five-fold cross validation was conducted by default to obtain 5-fold CV *R*^2^ per gene per tissue. Only “significant” gene expression prediction models with 5-fold CV *R*^2^ > 0.005 were retained in the output files (see our explanation in the Method and Discussion sections). The estimated Bayesian cis-eQTL weights from these “significant” gene expression prediction models can be used to conduct TWAS using both individual-level and summary-level GWAS data and are shared with the public along with our TIGAR-V2 tool.

Further, we compared gene expression prediction models trained from GTEx V8 ^1,26,40^ by nonparametric Bayesian DPR method to the ones (i.e., PredictDB models, see Web Resources) trained from the same GTEx V8 reference data by Elastic-Net method using the PrediXcan tool.

#### Application TWAS of breast and ovarian cancer

We used TIGAR-V2 to conduct TWAS of breast and ovarian cancer by using the Bayesian cis-eQTL weights estimated from GTEx V8 ^10^ of breast mammary tissue and ovary tissue and summary-level GWAS data ^11,12^. The GWAS summary data of breast and ovarian cancer were respectively obtained from the Breast Cancer Association Consortium (BCAC) with 122,977 cases and 105,974 controls of European ancestry ^11^ and the Ovarian Cancer Association Consortium (OCAC) with 22,406 cases and 40,941 controls of European ancestry ^12^. We also compared with TWAS results using eQTL weights given by Elastic-Net method (i.e., PrediXcan), which were also generated by our TIGAR-V2 tool.

Analyses conducted in this study use de-identified transcriptomic and genetic data from GTEx V8 and summary-level GWAS data of breast and ovarian cancer, which are in accordance with the ethical standards of the Institutional Review Board (IRB) at Emory University.

## Results

### Bayesian DPR eQTL weights from GTEx V8

From the GTEx V8 reference data as described previously, a total of 1,104,305 “significant” gene expression prediction models with 5-fold CV *R*^2^ > 0.005 were successfully trained by TIGAR (using the nonparametric Bayesian DPR method), for genes on the autosomal chromosomes of 49 tissue types. The average and median number of gene expression prediction models obtained per tissue type was ~22.5K.The corresponding Bayesian DPR eQTL weights (i.e., effect sizes of cis-SNPs in the fitted gene expression prediction models by nonparametric Bayesian DPR method) are publicly available along with our TIGAR-V2 tool.

#### Model over-fitting due to small training sample sizes

We present the median 5-fold CV *R*^2^ and the median training *R*^2^ of genome-wide genes per tissue type by TIGAR in Figure 2B and 2C, respectively. Here, the 5-fold CV *R*^2^ approximates the prediction *R*^2^ in independent data. Surprisingly, we observed that larger median 5-fold CV *R*^2^ and training *R*^2^ values were obtained for tissue types with smaller sample size (Figure 2). For example, the top median 5-fold CV *R*^2^ values (~ 0.04) were obtained for kidney cortex tissue (cyan bar), various brain tissues (yellow bars), and uterus tissue (hot pink bar), which all have sample sizes ~ 100. Whereas tissues which have relatively large sample sizes 400 ~ 600 (muscle skeletal, skin, and whole blood) have median 5-fold CV *R*^2^ ≈ 0.02. This trend is further demonstrated in the density plots of 5-fold CV *R*^2^ and training *R*^2^ by TIGAR for all tissues with color coded with respect to their training sample sizes (Figure S1).

We suspect this controversial trend is mainly due to model over-fitting with small training sample sizes. To further investigate this, we take the gene expression prediction models fitted with breast (*N* = 337) and ovarian (*N* = 140) tissue types as examples. First, we down-sampled breast tissue samples to 140 to match with the sample size of ovarian tissue. Second, we trained both PrediXcan Elastic-Net and TIGAR nonparametric Bayesian DPR models on the down-sampled breast tissue data. Third, we made density plot of the 5-fold CV *R*^2^ and training *R*^2^ for genes that have 5-fold CV *R*^2^ greater than various threshold, (0.005, 0.01, 0.05, 0.1, 0.2) (Figures S2 and S3).

We found that the same over-fitting issue existed for both PrediXcan Elastic-Net and TIGAR DPR methods. That is, the down-sampled breast tissue with the same sample size 140 as the ovarian tissue showed similar density distributions with respect to training *R*^2^, which had larger median training *R*^2^ than the breast tissue with sample size 337. As for the 5-fold CV *R*^2^, genes with 5-fold CV *R*^2^ > 0.2 had similar distributions between down-sampled and original breast tissues (that is expected), whereas other groups of genes had similar distributions between down-sampled breast tissue and ovarian tissue that are of the same training sample size (that is controversial due to overfitting). We think this is mainly driven by genes with relatively small expression heritability that would require a larger sample size to ensure a less over-fitted model. Since TIGAR DPR method has higher power to fit gene expression precision models for genes with relatively small expression heritability, the TIGAR training results are affected more by this over-fitting issue.

#### Comparison with PrediXcan eQTL weights

Additionally, we compared the gene expression prediction model training results by TIGAR with the ones by PrediXcan using the same GTEx V8 reference data ^1,26,40^. From boxplots of medians (Figure S4) and density plots (Figure S5) of 5-fold CV *R*^2^ and training *R*^2^ by PrediXcan, we observed the similar overfitting trend –– relatively larger median 5-fold CV *R*^2^ and median training *R*^2^ values were obtained with relatively smaller training sample sizes. These findings are consistent with our TIGAR training results (Figure 2; Figure S1) as well as our down-sample investigation (Figures S2-3).

As shown in Figure S6, more consistent 5-fold CV *R*^2^ and training *R*^2^ were obtained by PrediXcan and TIGAR for genes that were of relatively larger sample sizes (yellowish colors) and relatively higher expression heritability. We found TIGAR had consistently better performance with fitting more valid gene expression prediction models for genes of relatively smaller expression heritability. In particular, the higher median 5-fold CV *R*^2^ shown in Figure S4 by PrediXcan is based on the group of valid genes with 5-fold CV *R*^2^ > 0.005 by PrediXcan that is only < 50% of the valid genes by TIGAR (Figure 3A; Figure S7). These findings are also consistent with previous studies ^3^.

**Figure 3.**
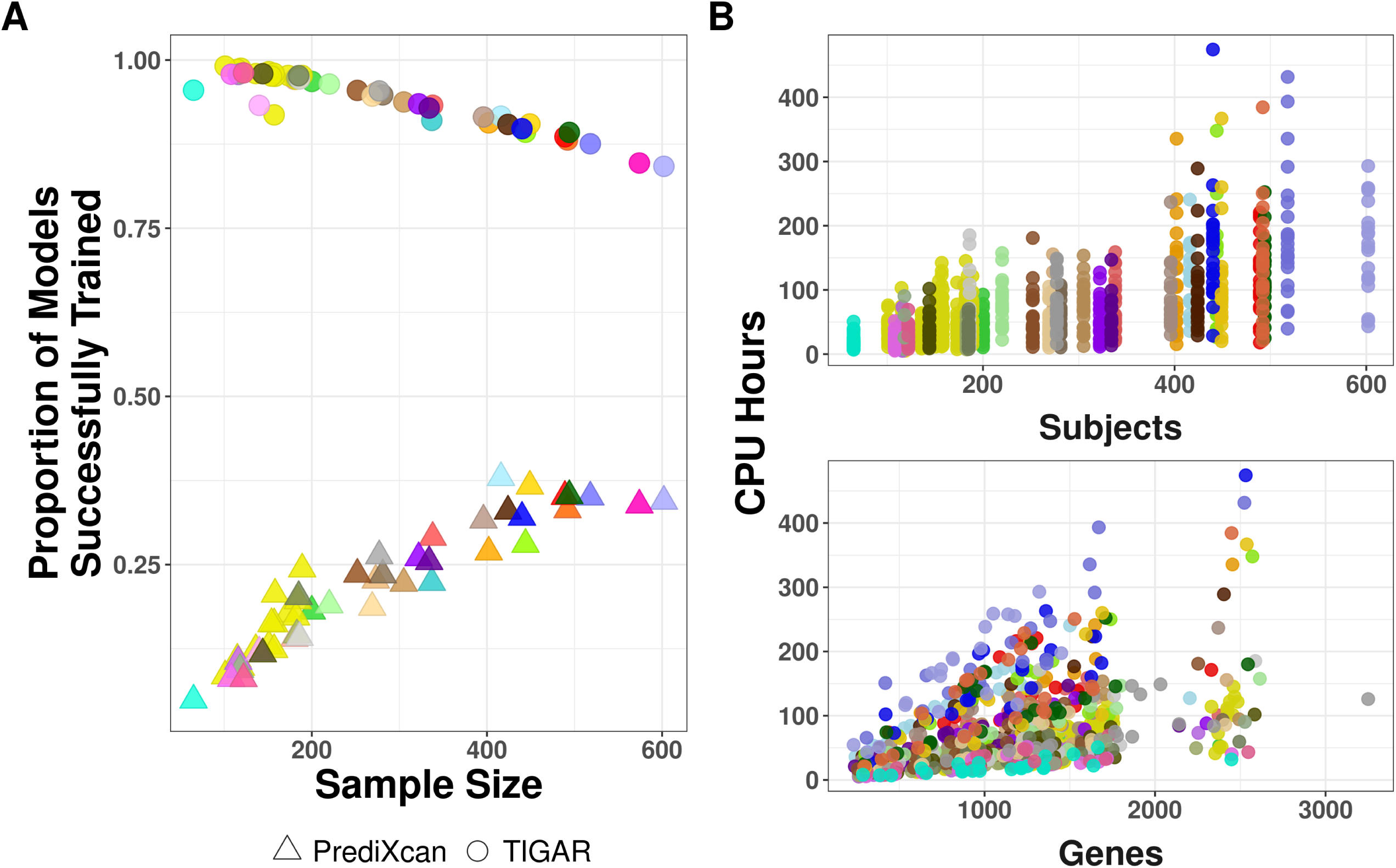
Proportion of valid gene expression prediction models by TIGAR vs. PrediXcan (A) and computation costs by TIGAR-V2 (B). The same color codes with respect to different tissue types as used in Figure 2 were used here. Computation times were in CPU Hours per chromosome per tissue for training gene expression prediction models with GTEx V8 reference data.

#### Computation cost by TIGAR-V2

The training computation costs in CPU hours per chromosome per tissue with GTEx V8 reference data by TIGAR-V2 are shown in Figure 3B, with respect to training sample sizes and number of genes in the chromosome. The computation cost per chromosome per tissue ranged from 5 CPU hours to over 474, with a median of 50.6 and mean of 69.1, which is mainly due to various numbers of genes per chromosome and various sample sizes per tissue. That is, with sample size ~ 300, the average computation time for training a nonparametric Bayesian DPR gene expression prediction model per gene with 5-fold cross-validation is only ~ 4 minutes by TIGAR-V2. The computation complexity is linear with respect to training sample sizes. Given the same computation cost for loading VCF genotype data, fitting a Bayesian DPR model costs about 2X computation time than fitting an Elastic-Net model by TIGAR-V2.

### TWAS of breast and ovarian cancer

From the gene expression prediction model training results by TIGAR, we respectively obtained 22,781 and 22,823 valid gene expression prediction models with 5-fold CV *R*^2^ > 0.005 by using the nonparametric Bayesian DPR method for breast (*N* = 337) and ovarian (*N* = 140) tissue types (Figure S7). Using GWAS summary statistics of breast cancer and ovarian cancer ^11,12^ and Bayesian *cis*-eQTL weights estimated with respect to the corresponding tissue type, TIGAR using our Bayesian eQTL weights respectively detected 88 and 37 significant TWAS genes (p-values < 2.5 × 10^−6^) for breast and ovarian cancer (see Manhattan plot in Figure 4). Of these significant genes, 17 were identified as risk genes of both breast and ovarian cancer (Table S1).

#### Independently significant TWAS risk genes by TIGAR

Out of these 88 significant TWAS genes for breast cancer by TIGAR, 20 genes are known GWAS risk genes of breast cancer ^11,41–48^, 64 are located within 1MB region of a previously identified GWAS loci of breast cancer ^11,41–48^ (Table S2), and 35 genes are identified by previous TWAS ^21,39,49–52^. Similarly, out of these 37 significant TWAS genes for ovarian cancer by TIGAR, 34 genes are located on chromosome 17 including two known GWAS risk genes (*NSF* and *PLEKHM1*) ^12,53,54^, 33 genes are located within 1MB of these two known GWAS risk genes (Table S3), and 13 genes (including *NSF* ^55^) are identified by previous TWAS ^39,55,56^. The known GWAS risk genes are curated from GWAS Catalogue ^57^ containing at least one significant SNP within or ±1MB around the gene region.

**Figure 4.**
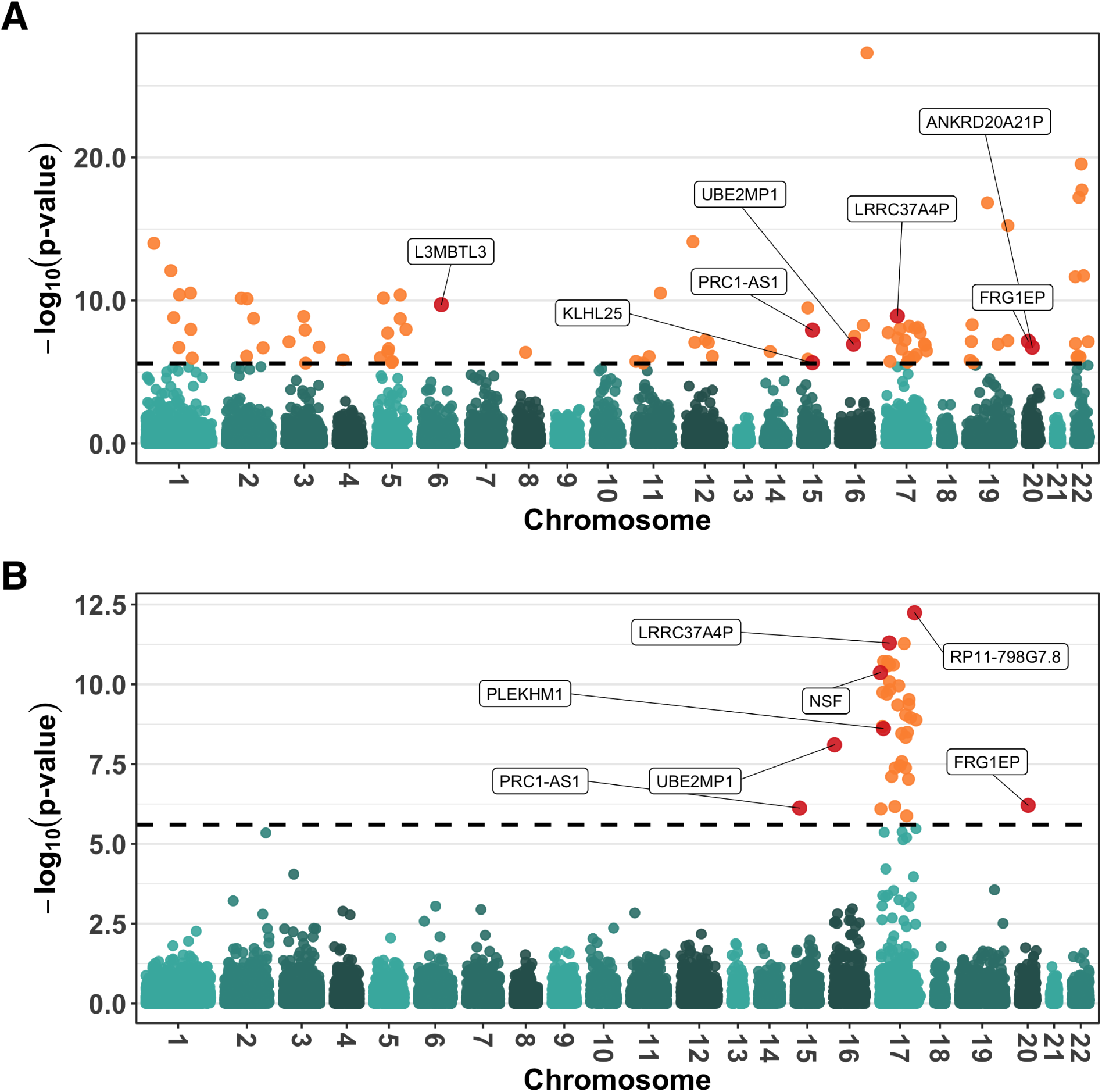
Manhattan plots of TWAS results by TIGAR for studying breast and ovarian cancer. (A) TWAS results of breast cancer with 88 significant risk genes. Significant gene *FCGR1B* of breast cancer (p-value: 4.12 × 10^−63^) was removed from (A) to reduce the upper limit of the y-axis. (B) TWAS results of ovarian cancer with 37 significant risk genes. Significant genes discussed in the main text are labeled in the plots.

Since TWAS is conducted using genotype data within ±1MB region of the test gene (i.e., test region), genes with overlapped test regions often have highly correlated GReX values (see locus-zoom plots around the top significant TWAS genes on chromosome 17 for breast and ovarian cancer in Figure 5). Thus, these nearby significant TWAS genes are often not representing independent associations. In Tables 1 and 2, we listed the most significant genes among genes that have shared test regions, which represent the independently significant TWAS risk genes. For breast cancer, 31 out of all 34 independent TWAS risk genes were either identified by previous GWAS/TWAS or within ±1MB region of previously identified risk genes of breast cancer (Table 1). For example, TIGAR identified *L3MBTL3* (previously identified by GWAS ^11^ and TWAS ^21,50–52^) and an additional 6 significant genes within the 1MB region of *L3MBTL3*. Of the independent TWAS genes of breast cancer, 17 (54%) have been identified by previous TWAS using PrediXcan and FUSION ^21,39,49–52^.

**Table 1.**
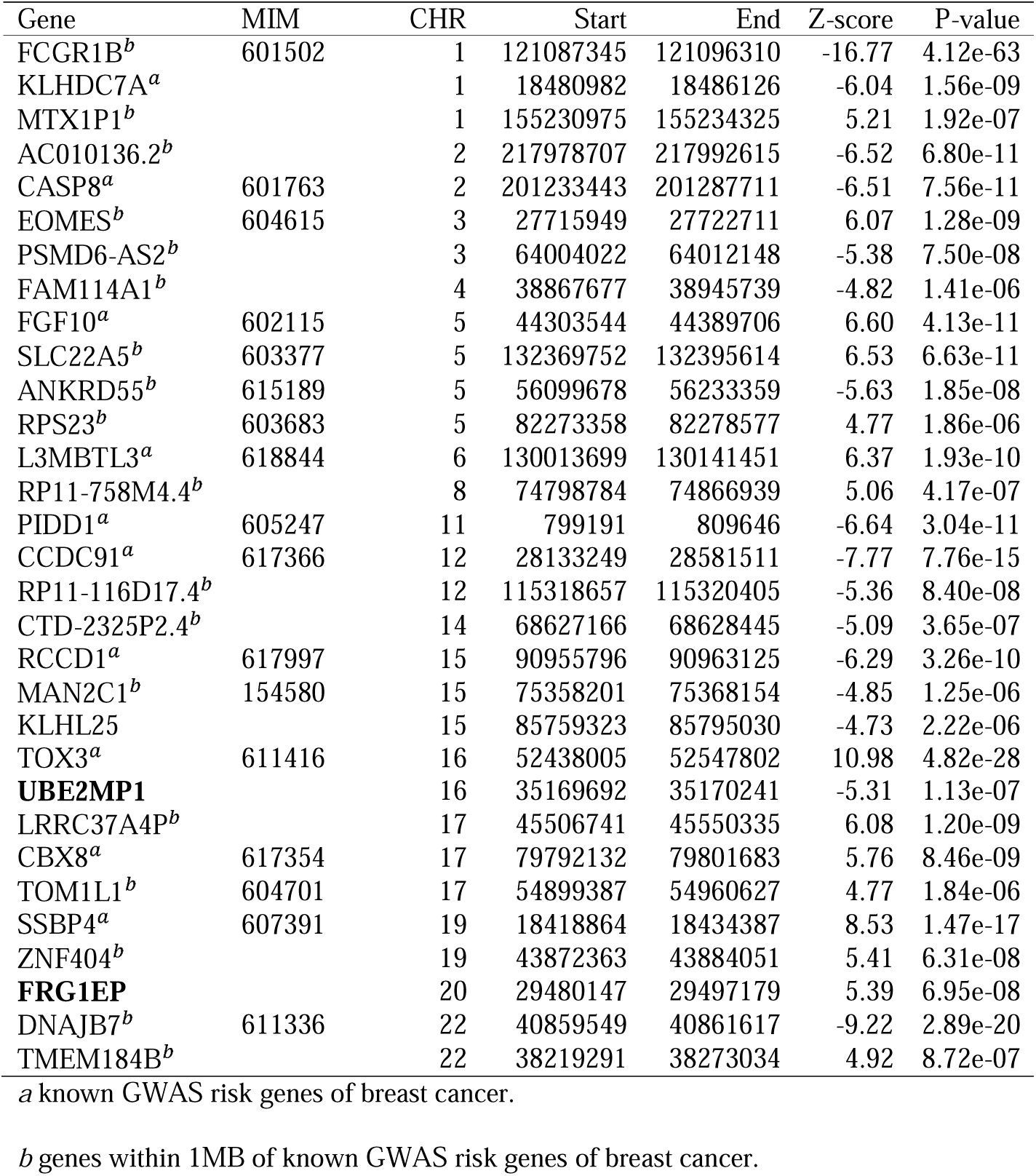
Independent TWAS risk genes of breast cancer identified by TIGAR, with novel risk genes of bold text.

| Gene | MIM | CHR | Start | End | Z-score | P-value |
| --- | --- | --- | --- | --- | --- | --- |
| FCGR1B<sup>b</sup> | 601502 | 1 | 121087345 | 121096310 | -16.77 | 4.12e-63 |
| KLHDC7A<sup>a</sup> |  | 1 | 18480982 | 18486126 | -6.04 | 1.56e-09 |
| MTX1P1<sup>b</sup> |  | 1 | 155230975 | 155234325 | 5.21 | 1.92e-07 |
| AC010136.2<sup>b</sup> |  | 2 | 217978707 | 217992615 | -6.52 | 6.80e-11 |
| CASP8<sup>a</sup> | 601763 | 2 | 201233443 | 201287711 | -6.51 | 7.56e-11 |
| EOMES<sup>b</sup> | 604615 | 3 | 27715949 | 27722711 | 6.07 | 1.28e-09 |
| PSMD6-AS2<sup>b</sup> |  | 3 | 64004022 | 64012148 | -5.38 | 7.50e-08 |
| FAM114A1<sup>b</sup> |  | 4 | 38867677 | 38945739 | -4.82 | 1.41e-06 |
| FGF10<sup>a</sup> | 602115 | 5 | 44303544 | 44389706 | 6.60 | 4.13e-11 |
| SLC22A5<sup>b</sup> | 603377 | 5 | 132369752 | 132395614 | 6.53 | 6.63e-11 |
| ANKRD55<sup>b</sup> | 615189 | 5 | 56099678 | 56233359 | -5.63 | 1.85e-08 |
| RPS23<sup>b</sup> | 603683 | 5 | 82273358 | 82278577 | 4.77 | 1.86e-06 |
| L3MBTL3<sup>a</sup> | 618844 | 6 | 130013699 | 130141451 | 6.37 | 1.93e-10 |
| RP11-758M4.4<sup>b</sup> |  | 8 | 74798784 | 74866939 | 5.06 | 4.17e-07 |
| PIDD1<sup>a</sup> | 605247 | 11 | 799191 | 809646 | -6.64 | 3.04e-11 |
| CCDC91<sup>a</sup> | 617366 | 12 | 28133249 | 28581511 | -7.77 | 7.76e-15 |
| RP11-116D17.4<sup>b</sup> |  | 12 | 115318657 | 115320405 | -5.36 | 8.40e-08 |
| CTD-2325P2.4<sup>b</sup> |  | 14 | 68627166 | 68628445 | -5.09 | 3.65e-07 |
| RCCD1<sup>a</sup> | 617997 | 15 | 90955796 | 90963125 | -6.29 | 3.26e-10 |
| MAN2C1<sup>b</sup> | 154580 | 15 | 75358201 | 75368154 | -4.85 | 1.25e-06 |
| KLHL25 |  | 15 | 85759323 | 85795030 | -4.73 | 2.22e-06 |
| TOX3<sup>a</sup> | 611416 | 16 | 52438005 | 52547802 | 10.98 | 4.82e-28 |
| **UBE2MP1** |  | 16 | 35169692 | 35170241 | -5.31 | 1.13e-07 |
| LRRC37A4P<sup>b</sup> |  | 17 | 45506741 | 45550335 | 6.08 | 1.20e-09 |
| CBX8<sup>a</sup> | 617354 | 17 | 79792132 | 79801683 | 5.76 | 8.46e-09 |
| TOM1L1<sup>b</sup> | 604701 | 17 | 54899387 | 54960627 | 4.77 | 1.84e-06 |
| SSBP4<sup>a</sup> | 607391 | 19 | 18418864 | 18434387 | 8.53 | 1.47e-17 |
| ZNF404<sup>b</sup> |  | 19 | 43872363 | 43884051 | 5.41 | 6.31e-08 |
| **FRG1EP** |  | 20 | 29480147 | 29497179 | 5.39 | 6.95e-08 |
| DNAJB7<sup>b</sup> | 611336 | 22 | 40859549 | 40861617 | -9.22 | 2.89e-20 |
| TMEM184B<sup>b</sup> |  | 22 | 38219291 | 38273034 | 4.92 | 8.72e-07 |

<sup>a</sup> known GWAS risk genes of breast cancer.
<sup>b</sup> genes within 1MB of known GWAS risk genes of breast cancer.

**Table 2.**
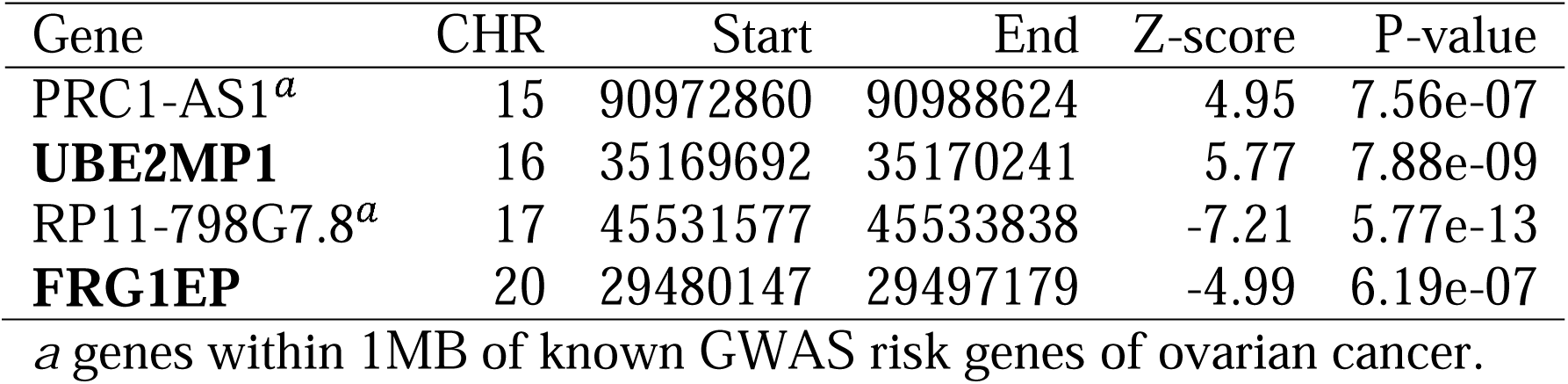
Independent TWAS risk genes of ovarian cancer identified by TIGAR, with novel risk genes of bold text. No MIM identifier available for any gene in this table.

| Gene | CHR | Start | End | Z-score | P-value |
| --- | --- | --- | --- | --- | --- |
| PRC1-AS1<sup>a</sup> | 15 | 90972860 | 90988624 | 4.95 | 7.56e-07 |
| **UBE2MP1** | 16 | 35169692 | 35170241 | 5.77 | 7.88e-09 |
| RP11-798G7.8<sup>a</sup> | 17 | 45531577 | 45533838 | -7.21 | 5.77e-13 |
| **FRG1EP** | 20 | 29480147 | 29497179 | -4.99 | 6.19e-07 |

<sup>a</sup> genes within 1MB of known GWAS risk genes of ovarian cancer.

**Figure 5.**
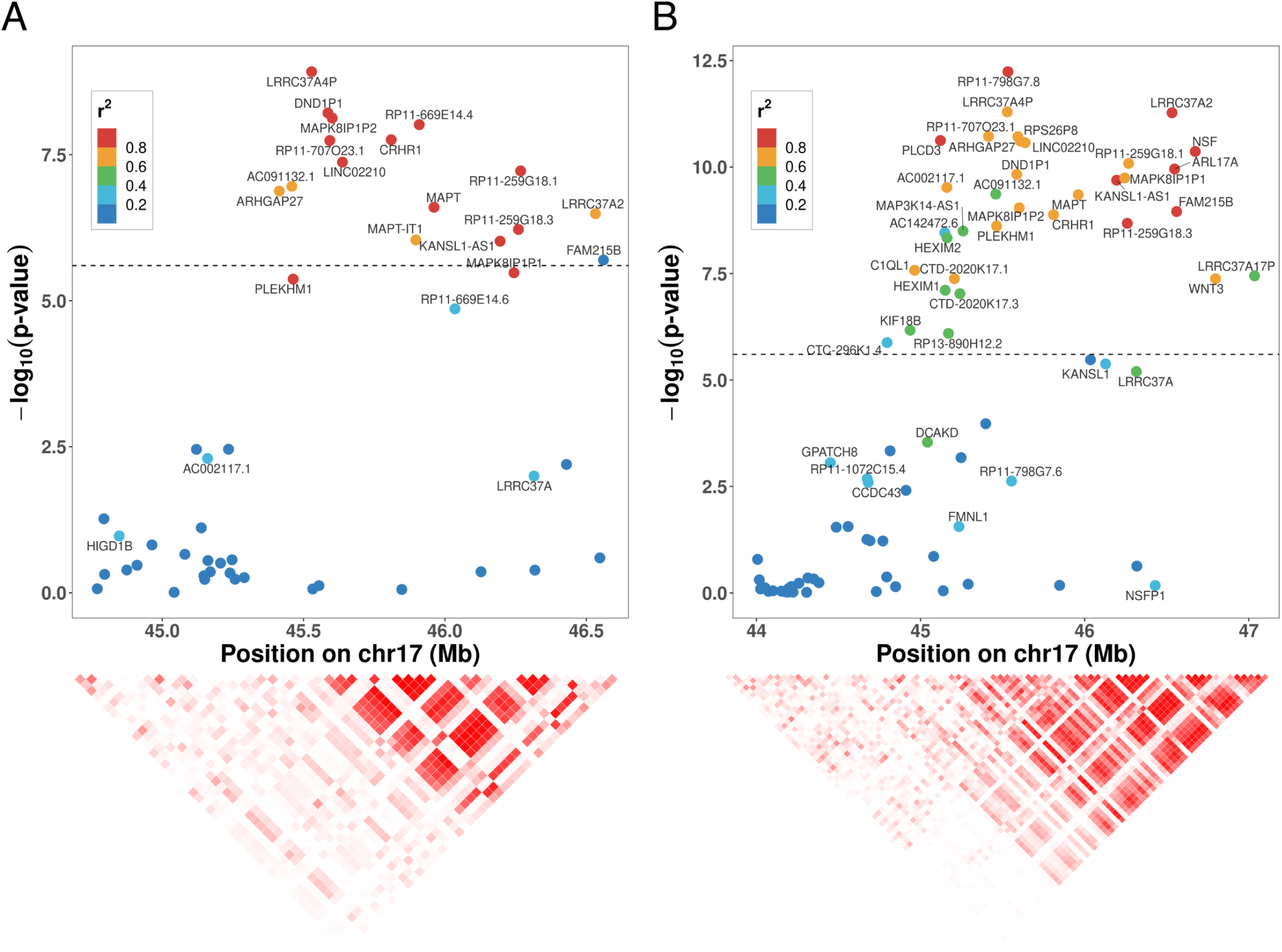
LocusZoom plots for genes within 1MB around the most significant TWAS gene on chromosome 17 of breast (A: *LRRC37A4P*) and ovarian (B: *RP11-789G7*.*8*) cancer. Each dot denotes the −lo*g*10(TWAS p-value) of a gene with color coded with respect to their GReX *R*^2^ with the top significant gene. The bottom heatmap colors denote the pairwise GReX *R*^2^ with bright red denoting GReX *R*^2^ close to 1 and white denoting GReX *R*^2^ close to 0.

Similarly, as shown in Table 2, TIGAR identified 4 independent significant TWAS genes for ovarian cancer. In particular, TWAS risk gene *RP11-798G7*.*8* on chromosome 17 was identified by previous TWAS ^39^ and lies within 1MB of known GWAS risk gene *PLEKHM1* ^12,53^. Interestingly, all independent TWAS risk genes of ovarian cancer by TIGAR (*PRC1-AS1, UBE2MP1, RP11-798G7*.*8*, and *FRG1EP*) are also TWAS risk genes of breast cancer ^11,39,58^, which demonstrates likely pleiotropy effect for these TWAS risk genes.

#### Significant TWAS risk genes identified by TIGAR in the 17q21.31 region

In particular, for the cluster of TWAS significant genes on chromosome 17 that were found to be shared by both breast and ovarian cancer, these genes have highly correlated GReX values as shown in Figure 5, including corticotrophin releasing hormone receptor 1 (*CRHR1*) and microtubule-associated protein tau (*MAPT*). These genes are located in the 17q21.31 region which contains a common inversion polymorphism of approximately 900KB in populations with European ancestry^59,60^, where two divergent *MAPT* haplotypes, H1 and H2, are shown to be associated with neurodegenerative diseases. A recent study showed that the expression of several genes in and at the borders of the inversion to be affected by the inversion; specific either to whole blood or different regions of the human brain^61^. Our findings show that the clusters of TWAS significant genes in the 17q21.31 region have differential GReX values in breast and ovary tissues with respect to both breast and ovarian cancers, and these GReX values are regulated by cis-eQTL part of the inversion polymorphism.

#### Novel findings by TIGAR

TIGAR identified 3 novel independent TWAS risk genes (*KLHL25, UBE2MP1*, and *FRG1EP*) for breast cancer. Gene *KLHL25* has known biological functions involved in carcinogenesis, while genes *UBE2MP1* and *FRG1EP* are near such a gene ^62–66^. Interestingly, genes *UBE2MP1* and *FRG1EP* were also identified for ovarian cancer by TIGAR (Table 2), and all three genes are involved with biological functions in carcinogenesis, either directly or indirectly. The protein encoded by *KLHL25* was reported acting as an adaptor protein for a suspected lung cancer tumour-suppressing protein *CUL3* to form an enzyme complex that targets ACLY, a protein often over-expressed in cancers, for degradation ^64^. Pseudogene *UBE2MP1* was found to have a significant expression-methylation-correlation difference between normal and cancerous breast tissue ^62^. *UBE2MP1* was also found to be amplified in gastric cancers [MIM: 613659] with amplified copy number variations in the 16p11.2 region, a mutation found to be associated with shorter overall survival ^66^, and was predicted to be a driver of lung adenocarcinoma [MIM: 211980] ^65^. The test region of *FRG1EP* overlaps with the test region of pseudogene *ANKRD20A21P*, another TWAS risk gene identified by TIGAR, which has been implicated as a potentially important lncRNA regulator of endometrial carcinogenesis [MIM: 608089] ^63^.

#### Comparison with TWAS results by PrediXcan

Additionally, we compared with the TWAS results of breast and ovarian cancer obtained by using *cis*-eQTL weights estimated by Elastic-Net method as used by PrediXcan (Tables S4 and S5), which were generated using the PrediXcan (Elastic-Net) function enabled in our TIGAR-V2 tool. We respectively obtained 11,095 and 12,337 valid gene expression prediction models for breast and ovary tissue types by using the PrediXcan function, about half of the number of valid gene expression prediction models by TIGAR (Figure S7). This is consistent with our above comparison of trained valid gene expression models by TIGAR and PrediXcan (Figure 3A).

As a result, PrediXcan detected 56 significant (32 independent) TWAS genes for breast cancer and 4 significant (2 independent) TWAS genes for ovarian cancer (Figure S8, Tables S4 and S5). Respective 30 out of 32 and 2 out of 2 of the independent TWAS risk genes by PrediXcan for breast and ovarian cancer were either identified by previous corresponding GWAS or within 1MB region of a known GWAS risk gene (Tables S6 and S7).

Even though there were respective 18 (56.25%) and 2 (100%) independent TWAS genes of breast and ovarian cancer by PrediXcan also identified by TIGAR (see Figure S9 and Tables S8, S9, S10), only TIGAR identified the novel TWAS genes *UBE2MP1* and *FRG1EP* shared by both breast and ovarian cancer and the known GWAS risk genes *FGF10* ^11,67^ and *TOX3* ^43,45^ of breast cancer. Other exclusive independent TWAS genes identified by TIGAR include lncRNA *RP11-758M4*.*4* which was shown to be a potential biomarker of breast cancer ^68^, *RPS23* which was found to be over-expressed in advanced colorectal adenocarcinomas [MIM: 114500] ^69^, and *ZNF404* whose dysregulation was linked to breast cancer pathogenesis by eQTL analyses ^70,71^. Potentially novel TWAS risk genes by PrediXcan and TIGAR that were not identified by previous GWAS are presented in Table S11.

#### Cis-eQTL weights by PrediXcan and TIGAR

To investigate the reason of why PrediXcan and TIGAR lead to different TWAS findings, we took three TWAS risk genes shared by both breast and ovarian cancer as examples. In particular, *FRG1EP* was only identified by TIGAR for both breast and ovarian cancer, while *LRRC37A4P* and *PRC1-AS1* were identified by both PrediXcan and TIGAR for both breast and ovarian cancer. Pseudogene *LRRC37A4P* on chromosome 17 lies within 1MB downstream from the known risk gene *PLEKHM1* of breast cancer and ovarian cancer ^12,53^. Gene *PRC1-AS1* on chromosome 15 is a long non-coding RNA (lncRNA) gene previously identified as being associated with breast carcinoma ^11,58^. Regulation of *PRC1-AS1* is known to differ with respect to different types of breast cancers ^72^ and increased expression of *PRC1-AS1* lncRNA is associated with hepatocellular carcinoma [MIM: 114550] ^73^.

We plotted the cis-eQTL weights estimated by Elastic-Net (PrediXcan) and Bayesian DPR method (TIGAR) from GTEx V8 for these three example TWAS risk genes, with color coded with respect to −*log*10(p-value) by single variant GWAS (Figures S10-S12). We observed that Bayesian estimates generally had non-zero values for all SNPs within the test region while Elastic-Net estimates had non-zero values for < 100 SNPs within the test region that had effect sizes (i.e., weights) of relatively larger magnitudes. These results match with the assumptions by the nonparametric Bayesian DPR (TIGAR) and Elastic-Net methods (PrediXcan). We can see that PrediXcan would miss the risk gene if test SNPs with non-zero weights have nonsignificant GWAS p-values such as *FRG1EP* (Figure S10). Otherwise, both PrediXcan and TIGAR would have similar power to identify the risk genes such as *LRRC37A4P* and *PRC1-AS1*, whose TWAS association are mainly driven by GWAS significant SNPs (Figures S11 and S12).

## Discussion

In this work, we develop a new version of the TIGAR tool ^3^ with improved computation efficiency, referred to as TIGAR-V2. Compared to the initial TIGAR tool, this new version reduces up to 90% computation time and up to 50% memory usage, mainly due to improved genotype data loading from VCF files and the usage of the Python library “numpy”. TIGAR-V2 can efficiently train gene expression prediction models by using both nonparametric Bayesian DPR and Elastic-Net (as used by PrediXcan) methods, as well as construct gene-based association tests using either individual-level or summary-level GWAS data. Gene-based associated tests implemented in TIGAR-V2 include both Burden statistics (based on FUSION ^2^ and S-PrediXcan ^26^ Z-score test statistics) and Variance-Component statistics.

We trained gene expression prediction models of 49 tissue types with the GTEx V8 reference data by using the nonparametric Bayesian DPR method. We provide trained eQTL weights of genes that have 5-fold CV *R*^2^ > 0.005 in the Synapse data base with a link given in Web Resources of in this paper. Since we used a more liberal threshold than the 0.01 used by previous studies ^2,21,22^ to allow more genes to be tested in follow-up TWAS, we would suggest users to investigate the 5-fold CV *R*^2^ and test p-values of the expression prediction models, as well as the biological functions of significant TWAS risk genes ^26^. Along with eQTL weights, we also provide gene information output files (an output file by TIGAR-V2) containing gene annotations (position, ID, name), training sample sizes, numbers of considered cis-SNPs, numbers of effective cis-SNPs for follow-up TWAS with non-zero eQTL weights, 5-fold CV *R*^2^, training *R*^2^, and a test p-value with respect to training *R*^2^. These gene information output files can be used by users to investigate the model training metrics of their TWAS significant genes. A similar approach is also suggested by the recent TWAS paper using GTEx data by PrediXcan^26^, which does not filter out any genes but only investigates the test p-value with respect to the training *R*^2^ for significant TWAS genes. Additionally, a recent power analysis of TWAS suggested useful threshold of expression heritability > 0.04 for a causal model where gene expression is directly causal with respect to the phenotype, and a threshold of expression heritability > 0.06 for a pleiotropy model where true causal SNPs of the phenotype are also true causal eQTL with respect to gene expression ^74^, which allowed TWAS had higher power than single variant GWAS for a simulation cohort with sample size 2504 that was used as both training and test data. We would only suggest TWAS as a secondary analysis to standard single variant GWAS, instead of as a competing analysis. We want to remind the users that TWAS is essentially a gene-based association test that is not comparable to standard single variant GWAS, but TWAS can provide extra biological insights with respect to the transcriptome data.

We demonstrated the usefulness of these trained models by performing TWAS of breast and ovarian cancer by integrating the estimated cis-eQTL weights of relevant tissue types with the relevant GWAS summary statistics. Compared to the cis-eQTL weights estimated by PrediXcan with the GTEx V8 data and TWAS results by PrediXcan, our Bayesian cis-eQTL weights lead to not only a larger number of significant TWAS risk genes but also interesting novel TWAS risk genes with potential pleiotropy effects for breast and ovarian cancer. With a larger number of “valid” gene expression prediction model trained by the nonparametric Bayesian DPR method, TIGAR is expected to identify a larger number of TWAS risk genes than PrediXcan. Our TWAS results of breast and ovarian cancer validated our TIGAR-V2 tool with findings consistent with previous GWAS and TWAS, revealed biological insights for known GWAS risk genes (*NSF* and *PLEKHM1*) ^12,53,54^ in the 17q21.31 region on chromosome 17 with pleiotropy effects for both breast and ovarian cancer, and identified novel risk genes that were shown to be possibly involved in the biological mechanisms of oncogenesis.

TIGAR-V2 tool still has its limitations such as considering only cis-eQTL data and assuming a two-stage model for TWAS. There are many other alternative TWAS tools available to address these two limits. For example, BGW-TWAS ^4^ and MOSTWAS^22^ uses both cis- and trans-genotype data to train gene expression prediction model of the target gene, while CoMM ^75^ and PMR-Egger ^5,76^ assume a joint model with reference and test data that can achieve higher power when both data sets are homogeneous.

In conclusion, TIGAR-V2 tool along with Bayesian cis-eQTL weights and reference LD covariance data (European ancestry) estimated from the GTEx V8 reference data are freely shared with the public on GitHub and Synapse. Given the convenience of directly loading VCF genotype data saved per chromosome, flexibility of using different training models and TWAS test statistics, and efficient computation enabled by Python source code, we believe our improved TIGAR-V2 tool will provide a useful resource for mapping risk genes of complex diseases by TWAS.

## Supporting information

Suppemental note

## Supplemental Data

Supplemental data include 2 text sections describing the nonparametric Bayesian DPR model and derivations of the FUSION and S-PrediXcan Z-score test statistics, 11 tables, and 12 figures.

## Acknowledgements

RP and JY are supported by National Institutes of Health (NIH/NIGMS) grant award R35GM138313. MPE was supported by NIH/NIGMS grant award R01GM117946 and NIH/NIA grant award RF1AG071170.

## Declaration of Interests

The authors declare no competing interests.

### Data and Code Availability

- TIGAR-V2 tool generated in this study is available at GITHUB https://github.com/yanglab-emory/TIGAR.
- All Bayesian eQTL weights of 49 tissue types from GTEx V8 are available at Synapse https://www.synapse.org/TIGAR_V2_Resource_GTExV8.
- GTEx V8 data are available from dbGaP with accession phs000424.v8.p2.
- GWAS data for studying breast cancer is available from Breast Cancer Association Consortium (BCAC) http://bcac.ccge.medschl.cam.ac.uk/bcacdata/oncoarray/oncoarray-and-combined-summary-result/gwas-summary-results-breast-cancer-risk-2017/.
- GWAS data for studying ovarian cancer is available from Ovarian Cancer Association Consortium (OCAC) ftp://ftp.ebi.ac.uk/pub/databases/gwas/summary_statistics/PhelanCM_28346442_GCST004415/.

### Web Resources

- TIGAR-V2: https://github.com/yanglab-emory/TIGAR
- Synapse: https://www.synapse.org/TIGAR_V2_Resource_GTExV8
- GTEx portal: https://gtexportal.org/home/
- eQTL weights trained by PrediXcan with GTEx V8: https://predictdb.org
- OMIM: https://omim.org/

## Notes

https://www.synapse.org/TIGAR_V2_Resource_GTExV8

### Competing Interest Statement

The authors have declared no competing interest.

