## Supplementary material for "TIGAR-V2: Efficient TWAS Tool with Nonparametric Bayesian eQTL Weights of 49 Tissue Types from GTEx V8": Suppemental note

### Supplemental Figures

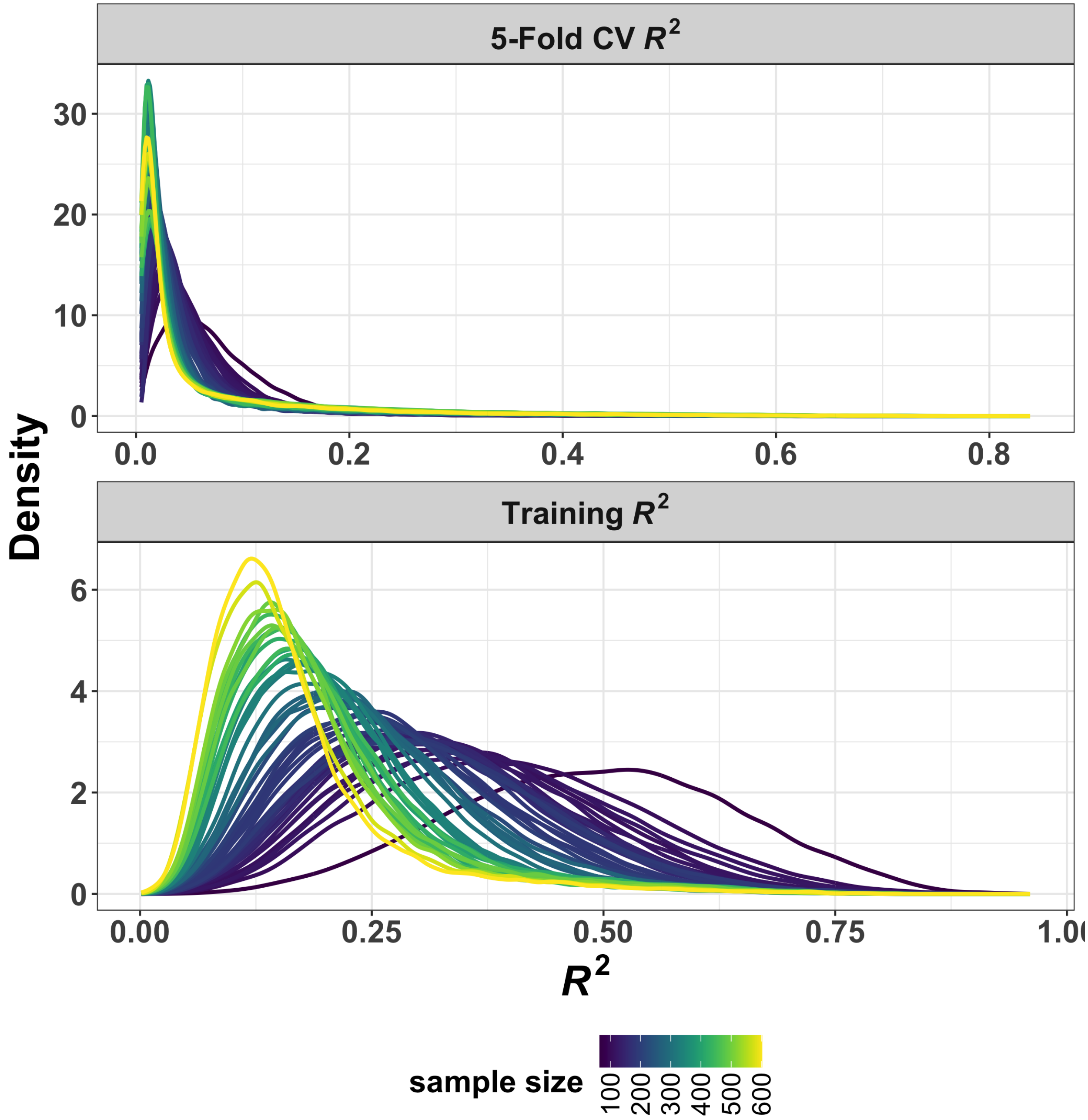


**Figure S1: Density Plots of 5-Fold CV $R^2$ and Training $R^2$ by TIGAR.** One line per tissue type with color coded according to training sample sizes.

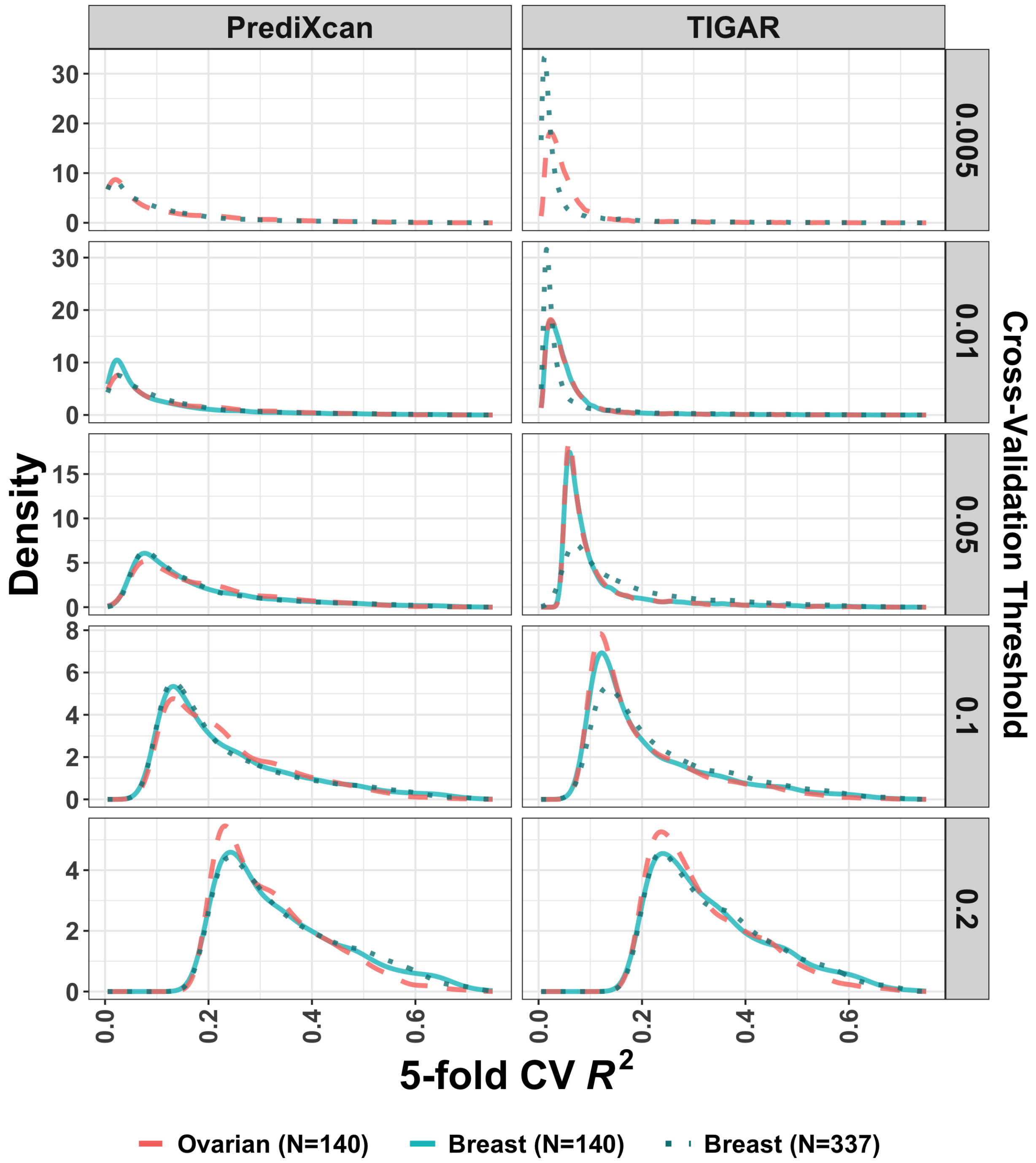


**Figure S2: Density plots of 5-fold CV $R^2$ by PrediXcan and TIGAR**. Training data of Ovarian ($N = 140$), Breast ($N = 337$), and downsampled Breast ($N = 140$) tissues were used. Density plots were stratified for groups of genes with 5-fold CV $R^2$ $>$ (0.005, 0.01, 0.05, 0.1, 0.2).

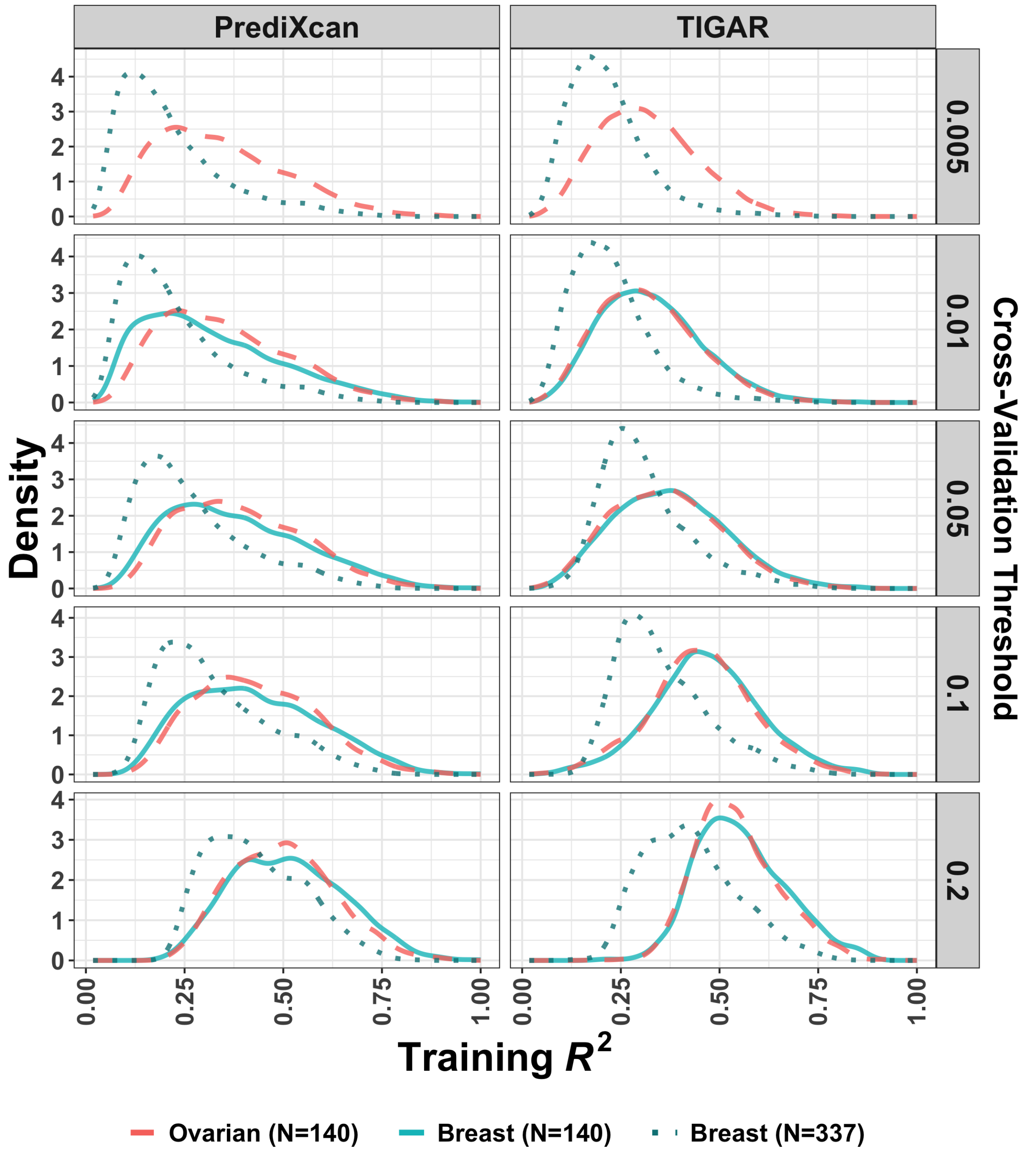


**Figure S3: Density plots of training $R^2$ by PrediXcan and TIGAR.** Training data of Ovarian ($N = 140$), Breast ($N = 337$), and downsampled Breast ($N = 140$) tissues were used. Density plots were stratified for groups of genes with 5-fold CV $R^2$ $>$ (0.005, 0.01, 0.05, 0.1, 0.2).

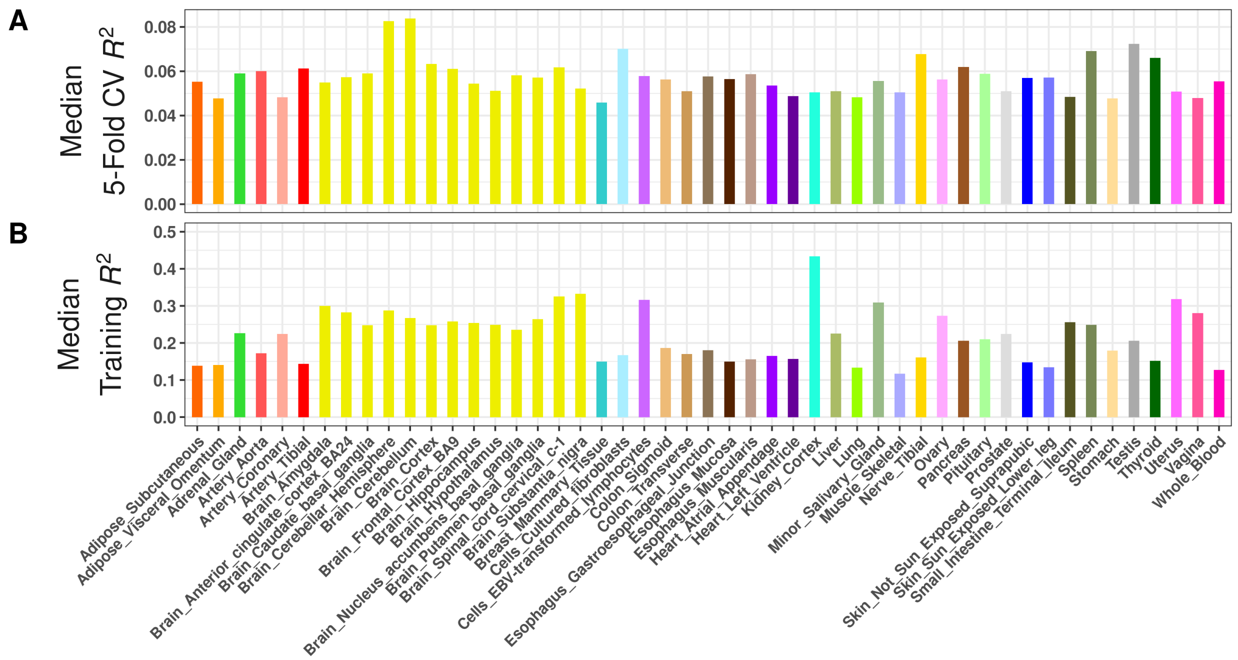


**Figure S4: Median 5-Fold CV $R^2$ (A) and Median Training $R^2$ (B) per tissue type for all trained gene expression prediction models with GTEx V8 reference data of 49 tissue types by PrediXcan.**

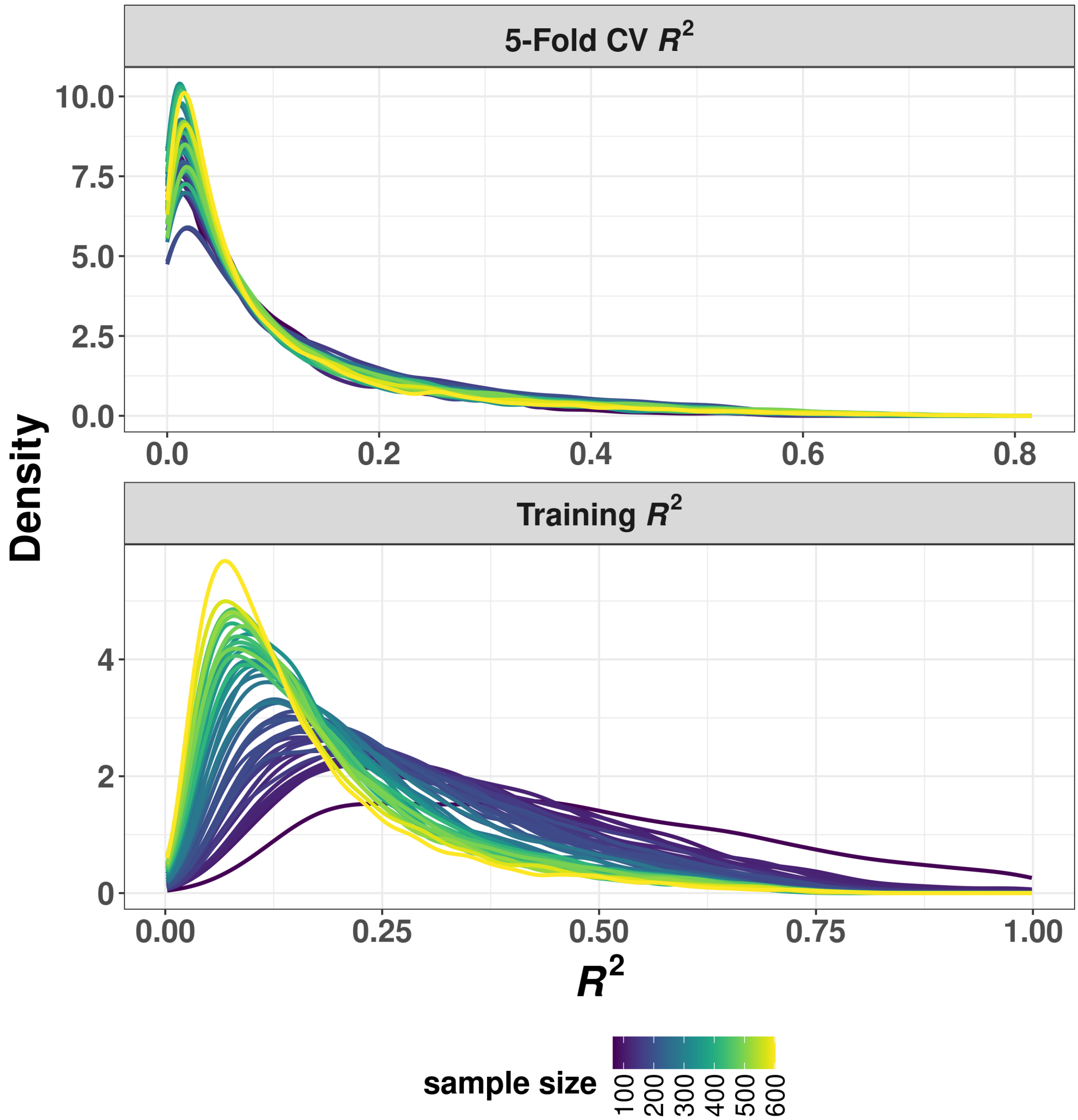


**Figure S5: Density Plots of 5-Fold CV $R^2$ and Training $R^2$ by PrediXcan for all tissue types of GTEx V8**. Color coded with respect to training sample sizes.

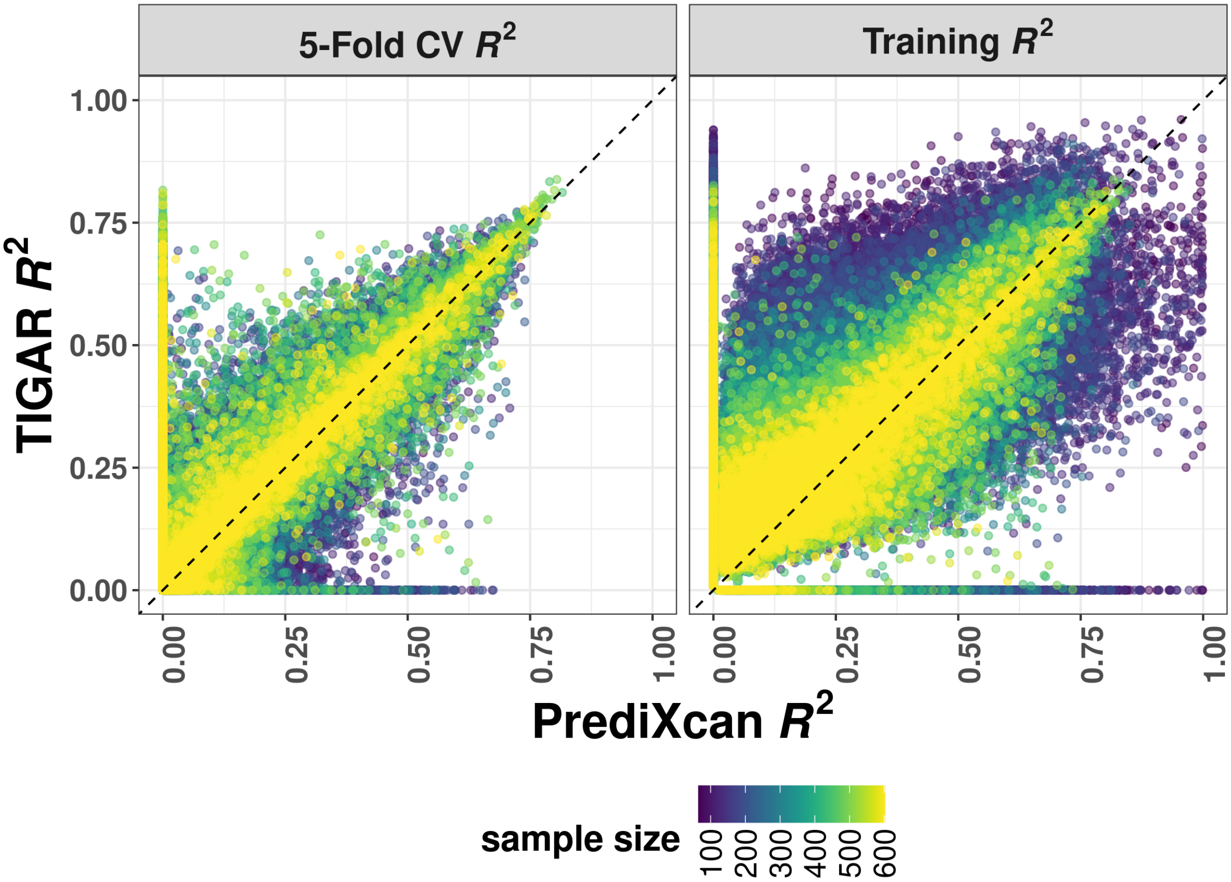


**Figure S6: Comparison of 5-Fold CV $R^2$ and Training $R^2$ by PrediXcan and TIGAR for all tissue types, colored by tissue sample size.**

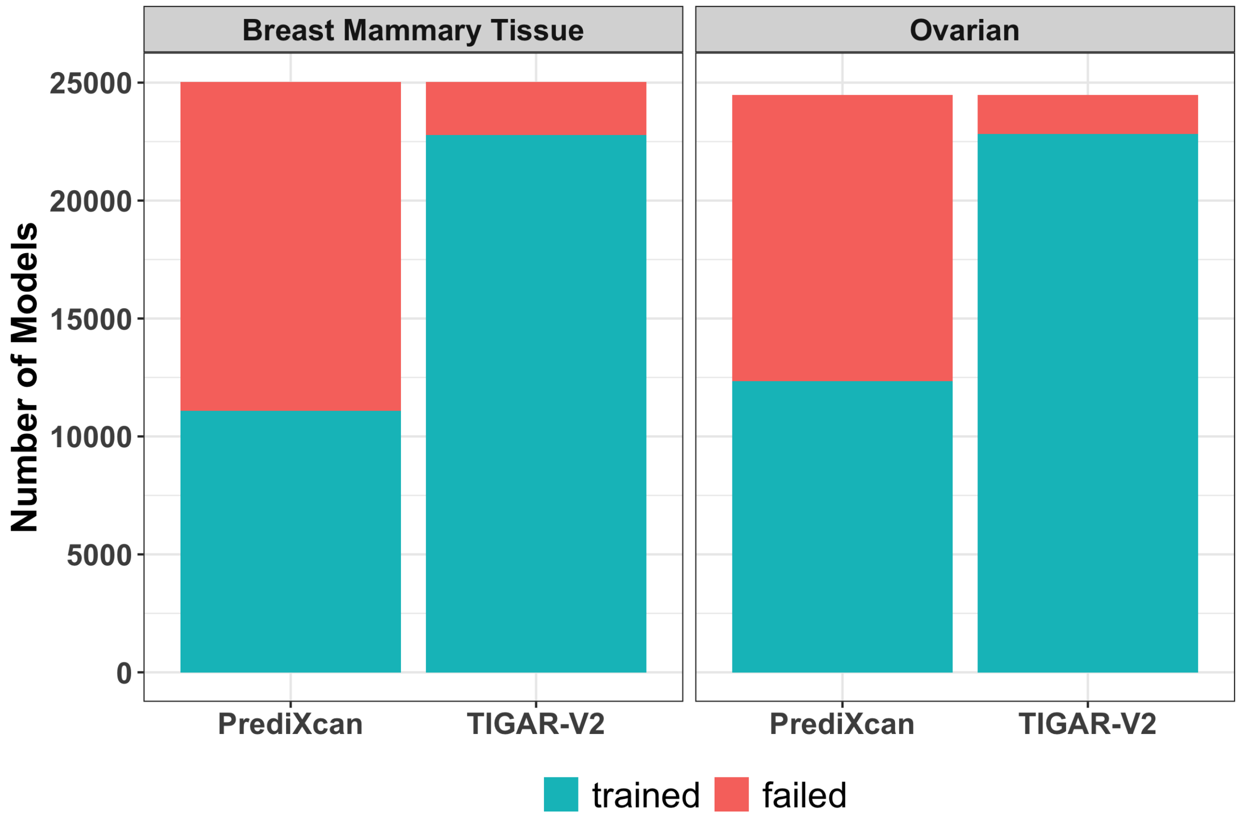


Figure S7: Number of failed and valid gene expression prediction models by PrediXcan and TIGAR for breast and ovarian tissues.

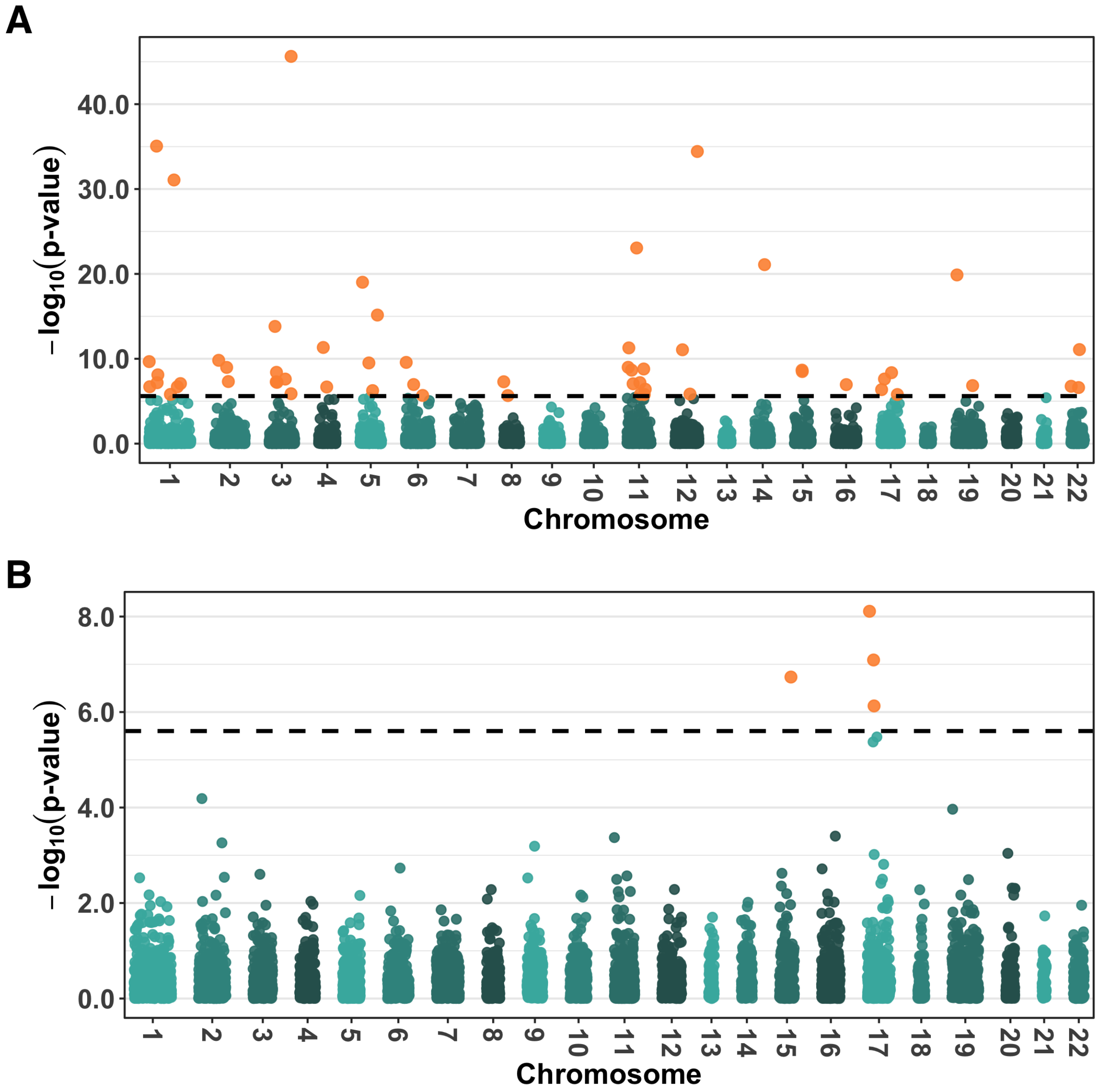


**Figure S8: Manhattan plots of TWAS results by PrediXcan.** (A) TWAS results for studying breast cancer with 56 significant genes; (B) TWAS results for studying ovarian cancer with 4 significant genes.

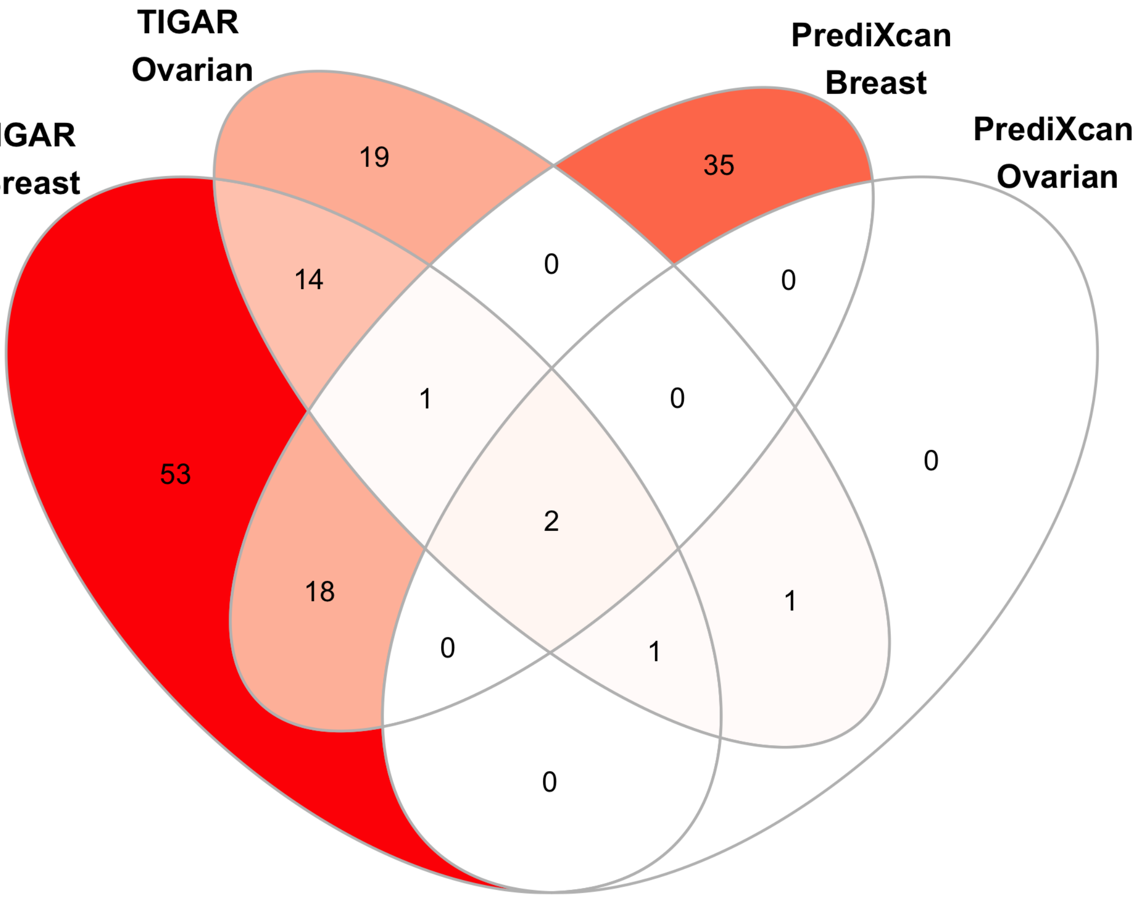


**Figure S9: Venn diagram of the number of shared TWAS risk genes by PrediXcan and TIGAR for breast and ovarian cancer.**

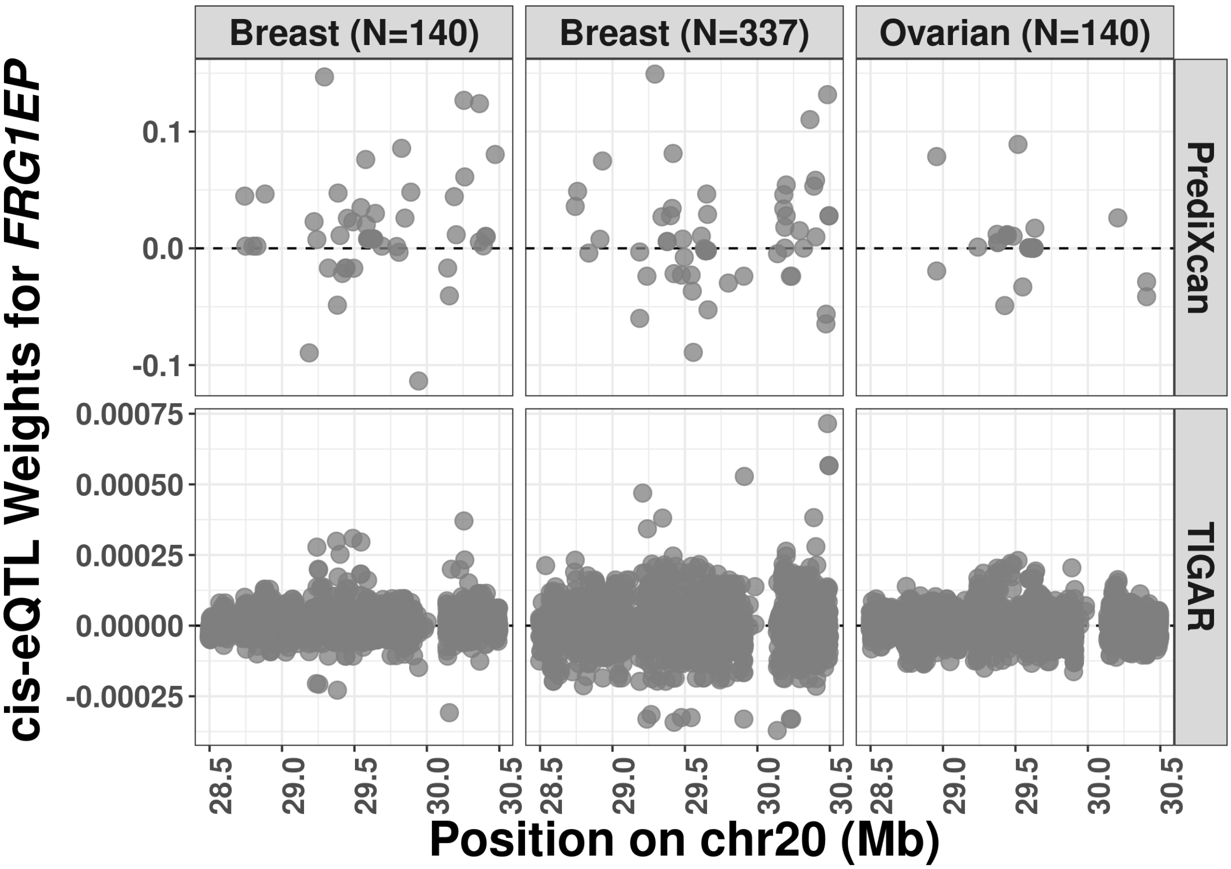


**Figure S10: Cis-eQTL weights estimated by PrediXcan and TIGAR for *FRG1EP* per training data set.** The novel TWAS risk gene *FRG1EP* was only identified by TIGAR for both breast and ovarian cancer. None of the SNPs has GWAS p-value $< 10^{-5}$.

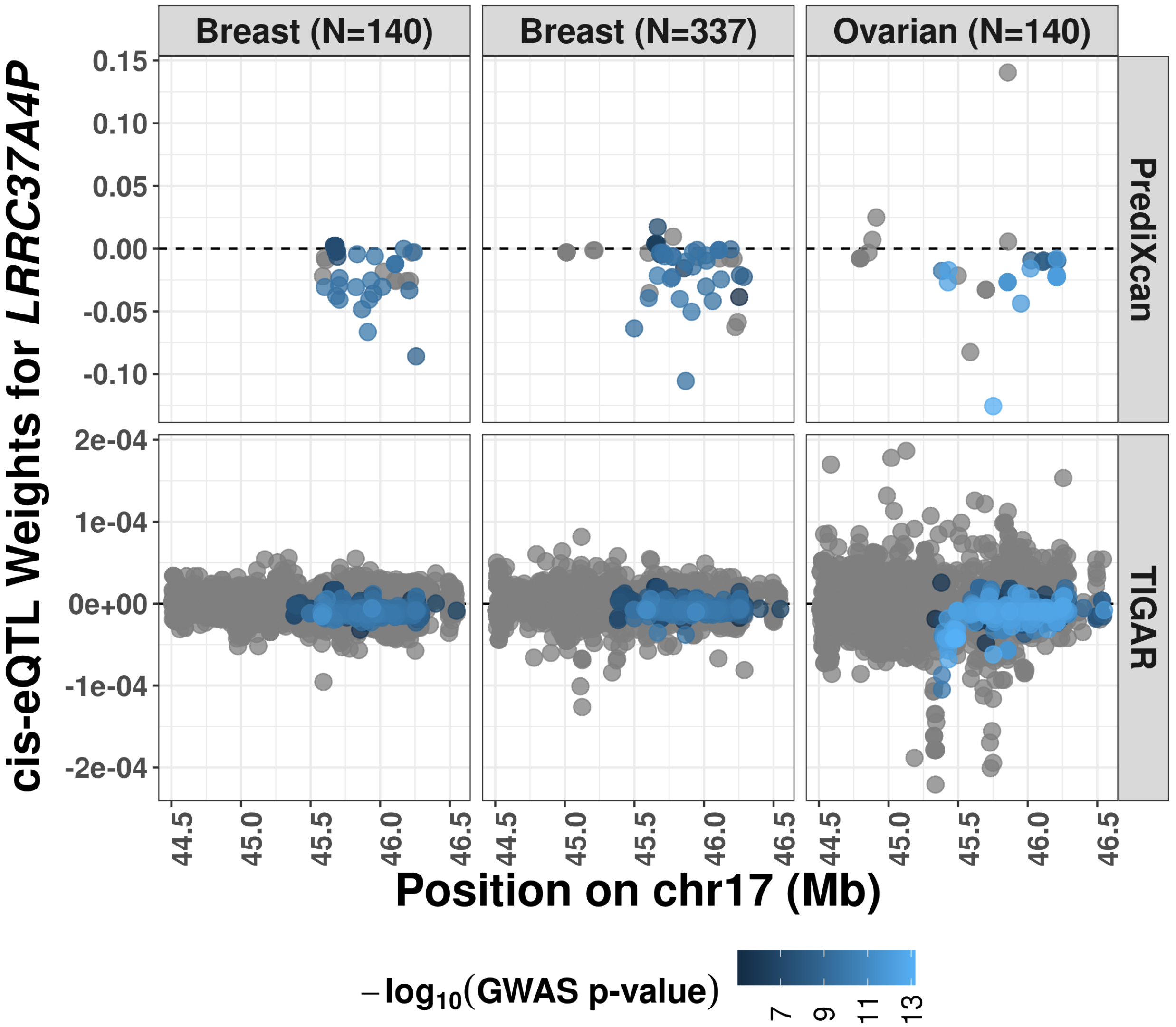


**Figure S11: Cis-eQTL weights estimated by PrediXcan and TIGAR for *LRRC37A4P* per training data set.** *LRRC37A4P* was identified by both PrediXcan and TIGAR for both breast and ovarian cancer.

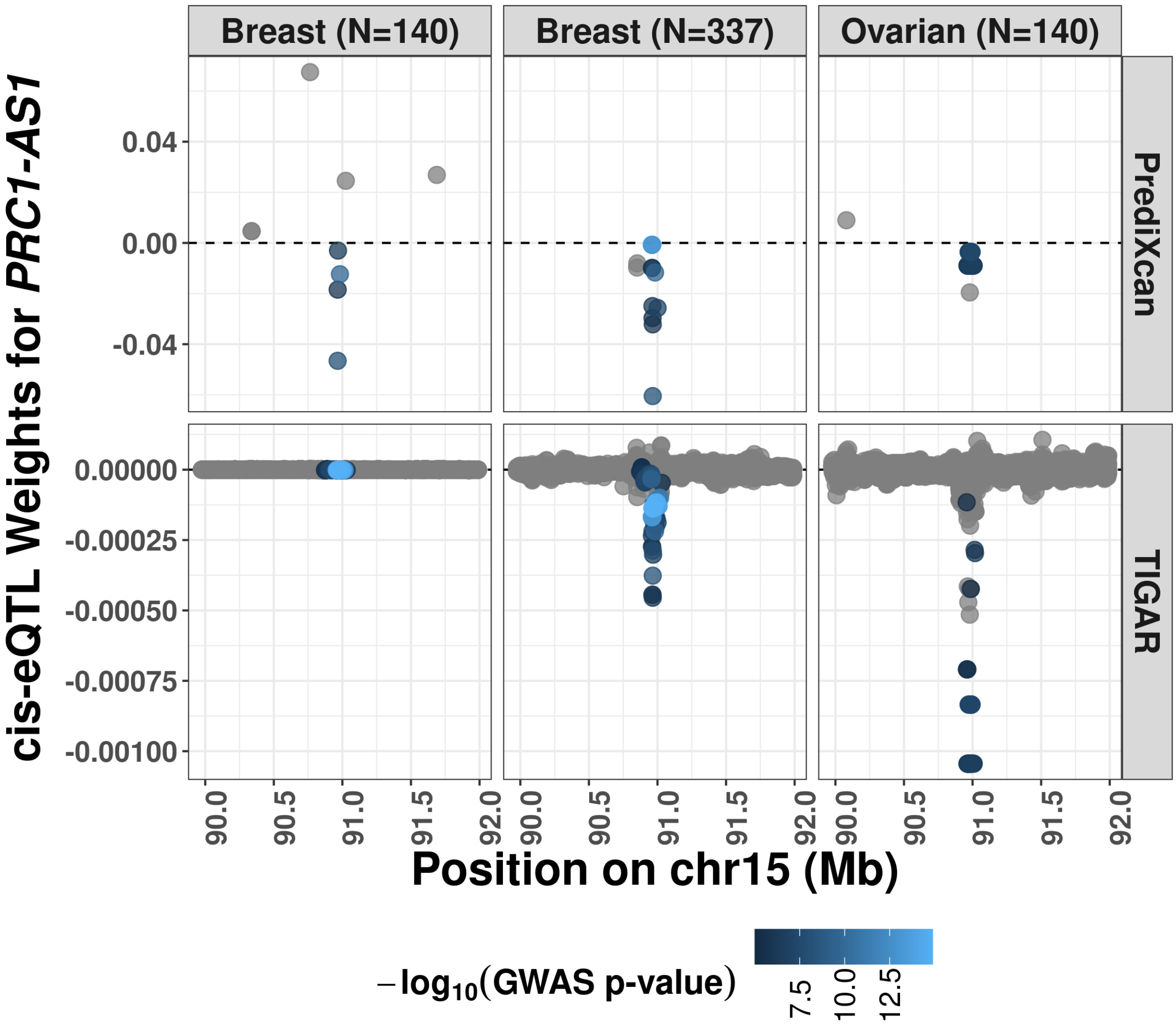


**Figure S12: Cis-eQTL weights by PrediXcan and TIGAR for *PRC1-AS1* per training data set**. *PRC1-AS1* was identified by both PrediXcan and TIGAR for both breast and ovarian cancer.

### Supplemental Tables

| Gene | MIM | CHR | Start | End | Breast | | Ovarian | |
|---|---|---|---|---|---|---|---|---|
| | | | | | Z | P-value | Z | P-value |
| PRC1-AS1[a,d] | | 15 | 90972860 | 90988624 | 5.70 | 1.17e-08 | 4.95 | 7.56e-07 |
| UBE2MP1 | | 16 | 35169692 | 35170241 | -5.31 | 1.13e-07 | 5.77 | 7.88e-09 |
| ARHGAP27[a,d] | 610591 | 17 | 45393902 | 45434421 | -5.27 | 1.33e-07 | 6.24 | 4.32e-10 |
| AC091132.1[b,d] | | 17 | 45452844 | 45464065 | -5.31 | 1.10e-07 | -6.36 | 2.04e-10 |
| LRRC37A4P[b,d] | | 17 | 45506741 | 45550335 | 6.08 | 1.20e-09 | 6.90 | 5.07e-12 |
| DND1P1[b,d] | | 17 | 45585871 | 45586929 | -5.81 | 6.11e-09 | -6.90 | 5.31e-12 |
| RP11-707O23.1[b,d] | | 17 | 45592621 | 45593369 | -5.63 | 1.81e-08 | -6.66 | 2.68e-11 |
| MAPK8IP1P2[b,d] | | 17 | 45600869 | 45602340 | -5.78 | 7.48e-09 | -6.71 | 1.91e-11 |
| LINC02210[b,d] | | 17 | 45620328 | 45655156 | -5.48 | 4.23e-08 | -6.50 | 8.23e-11 |
| CRHR1[b,d] | 122561 | 17 | 45784280 | 45835828 | -5.63 | 1.75e-08 | -6.71 | 1.93e-11 |
| MAPT[a,d] | 157140 | 17 | 45894382 | 46028334 | -5.16 | 2.51e-07 | 6.24 | 4.45e-10 |
| KANSL1-AS1[b,d] | | 17 | 46193576 | 46196723 | -4.90 | 9.57e-07 | -6.07 | 1.32e-09 |
| RP11-259G18.3[b,d] | | 17 | 46259551 | 46260606 | -4.99 | 6.03e-07 | -6.09 | 1.11e-09 |
| RP11-259G18.1[b,d] | | 17 | 46267037 | 46268694 | -5.42 | 5.99e-08 | -6.41 | 1.49e-10 |
| LRRC37A2[b,d] | 616556 | 17 | 46511511 | 46553449 | -5.11 | 3.24e-07 | -6.12 | 9.08e-10 |
| FAM215B[b,d] | | 17 | 46558830 | 46562795 | -4.75 | 2.01e-06 | -5.99 | 2.09e-09 |
| FRG1EP | | 20 | 29480147 | 29497179 | 5.39 | 6.95e-08 | -4.99 | 6.19e-07 |

*a* known GWAS risk gene of breast cancer.

*b* genes within 1MB of a known GWAS risk gene of breast cancer.

*c* genes within 1MB of a known GWAS risk gene of ovarian cancer.

**Table S1: Significant TWAS risk genes of both breast and ovarian cancer identified by TIGAR.**

| Gene | MIM | CHR | Start | End | Z-score | P-value |
|---|---|---|---|---|---|---|
| EMBP1 | | 1 | 121519112 | 121571892 | 7.16 | 8.13e-13 |
| KLHDC7A | | 1 | 18480982 | 18486126 | -6.04 | 1.56e-09 |
| ASH1L | 607999 | 1 | 155335268 | 155562807 | -4.88 | 1.05e-06 |
| CASP8 | 601763 | 2 | 201233443 | 201287711 | -6.51 | 7.56e-11 |
| SLC4A7 | 603353 | 3 | 27372721 | 27484420 | -5.71 | 1.14e-08 |
| NEK10 | 618726 | 3 | 27110085 | 27369460 | -5.22 | 1.79e-07 |
| FGF10 | 602115 | 5 | 44303544 | 44389706 | 6.60 | 4.13e-11 |
| MRPS30 | 611991 | 5 | 44808925 | 44820428 | -5.73 | 1.03e-08 |
| ATP6AP1L | | 5 | 82279462 | 82386977 | -4.74 | 2.15e-06 |
| L3MBTL3 | 618844 | 6 | 130013699 | 130141451 | 6.37 | 1.93e-10 |
| PIDD1 | 605247 | 11 | 799191 | 809646 | -6.64 | 3.04e-11 |
| CCDC91 | 617366 | 12 | 28133249 | 28581511 | -7.77 | 7.76e-15 |
| RCCD1 | 617997 | 15 | 90955796 | 90963125 | -6.29 | 3.26e-10 |
| PRC1-AS1 | | 15 | 90972860 | 90988624 | 5.70 | 1.17e-08 |
| TOX3 | 611416 | 16 | 52438005 | 52547802 | 10.98 | 4.82e-28 |
| CBX8 | 617354 | 17 | 79792132 | 79801683 | 5.76 | 8.46e-09 |
| ARHGAP27 | 610591 | 17 | 45393902 | 45434421 | -5.27 | 1.33e-07 |
| MAPT | 157140 | 17 | 45894382 | 46028334 | -5.16 | 2.51e-07 |
| SSBP4 | 607391 | 19 | 18418864 | 18434387 | 8.53 | 1.47e-17 |
| L3MBTL2 | 611865 | 22 | 41205205 | 41231271 | 8.63 | 6.03e-18 |

**Table S2: Known GWAS risk genes of breast cancer that were identified by TIGAR.**

| Gene | MIM | CHR | Start | End | Z-score | P-value |
|---|---|---|---|---|---|---|
| NSF | 601633 | 17 | 46590669 | 46757464 | 6.59 | 4.31e-11 |
| PLEKHM1 | 611466 | 17 | 45435900 | 45490749 | -5.97 | 2.43e-09 |

**Table S3: Known GWAS risk genes of ovarian cancer that were identified by TIGAR.**

| Gene | MIM | CHR | Start | End | Z-score | P-value |
|---|---|---|---|---|---|---|
| CH17-437K3.1[b] | | 1 | 121396754 | 121463129 | 12.49 | 8.75e-36 |
| ASH1L[a] | 607999 | 1 | 155335268 | 155562807 | -6.35 | 2.16e-10 |
| NSUN4[b] | 615394 | 1 | 46340177 | 46365152 | 5.77 | 7.81e-09 |
| KLHDC7A[a] | | 1 | 18480982 | 18486126 | -5.20 | 2.01e-07 |
| ALS2CR12[b] | | 2 | 201288271 | 201357398 | 6.40 | 1.52e-10 |
| RP11-337N6.3 | | 2 | 177317715 | 177318471 | 5.46 | 4.82e-08 |
| SLC4A7[a] | 603353 | 3 | 27372721 | 27484420 | -14.30 | 2.34e-46 |
| ZBTB38[a] | 612218 | 3 | 141324213 | 141449792 | -7.68 | 1.54e-14 |
| LINC00886[a] | | 3 | 156747346 | 156817062 | 5.89 | 3.96e-09 |
| CMSS1[a] | | 3 | 99817834 | 100181732 | 5.44 | 5.31e-08 |
| PSMD6-AS2[b] | | 3 | 64004022 | 64012148 | -5.42 | 5.99e-08 |
| EFCC1 | | 3 | 129001629 | 129040742 | 4.84 | 1.32e-06 |
| GLRA3[b] | 600421 | 4 | 174636914 | 174829314 | 6.92 | 4.66e-12 |
| PPM1K[b] | 611065 | 4 | 88257620 | 88284769 | -5.19 | 2.11e-07 |
| SLC22A5[b] | 603377 | 5 | 132369752 | 132395614 | 9.09 | 9.64e-20 |
| ANKRD55[b] | 615189 | 5 | 56099678 | 56233359 | -6.29 | 3.09e-10 |
| L3MBTL3[a] | 618844 | 6 | 130013699 | 130141451 | 6.32 | 2.67e-10 |
| TOB2P1[b] | | 6 | 28217643 | 28218634 | 5.31 | 1.09e-07 |
| ZNF703[b] | 617045 | 8 | 37695751 | 37700021 | 5.45 | 5.11e-08 |
| PRR33[b] | | 11 | 1888577 | 1891772 | -10.05 | 8.84e-24 |
| EFEMP2[b] | 604633 | 11 | 65866441 | 65873592 | 6.03 | 1.60e-09 |
| SPTY2D1[a] | | 11 | 18606401 | 18634791 | 4.80 | 1.57e-06 |
| NTN4[b] | 610401 | 12 | 95657807 | 95791152 | -12.37 | 3.75e-35 |
| RP11-967K21.1[b] | | 12 | 28163298 | 28190738 | 6.83 | 8.68e-12 |
| RPL12P7[b] | | 14 | 68693090 | 68693583 | 9.60 | 8.11e-22 |
| RCCD1[a] | 617997 | 15 | 90955796 | 90963125 | -5.98 | 2.18e-09 |
| RP11-212I21.2[b] | | 16 | 55426797 | 55462297 | 5.31 | 1.11e-07 |
| CBX8[a] | 617354 | 17 | 79792132 | 79801683 | 5.87 | 4.35e-09 |
| LRRC37A4P[b] | | 17 | 45506741 | 45550335 | 5.58 | 2.43e-08 |
| COX11[b] | 603648 | 17 | 54951902 | 54968764 | 4.79 | 1.63e-06 |
| LRRC25[b] | 607518 | 19 | 18391144 | 18397617 | 9.31 | 1.32e-20 |
| APOBEC3B[b] | 607110 | 22 | 38982347 | 38992804 | 6.83 | 8.25e-12 |

*a* a known GWAS risk gene of breast cancer.

*b* within 1MB of a known GWAS risk gene of breast cancer.

**Table S4: Independent TWAS risk genes of breast cancer identified by PrediXcan.**

| Gene | CHR | Start | End | Z-score | P-value |
|---|---|---|---|---|---|
| PRC1-AS1[a] | 15 | 90972860 | 90988624 | 5.21 | 1.85e-07 |
| LINC02210[a] | 17 | 45620328 | 45655156 | -5.77 | 7.76e-09 |

*a* within 1MB of a kown GWAS risk gene of ovarian cancer.

∗ No MIM identifier available for any gene in this table.

**Table S5: Independent TWAS risk genes of ovarian cancer identified by PrediXcan.**

| Gene | MIM | CHR | Start | End | Z-score | P-value |
|---|---|---|---|---|---|---|
| ASH1L | 607999 | 1 | 155335268 | 155562807 | -6.35 | 2.16e-10 |
| LINC00623 | | 1 | 120952439 | 121009291 | 5.40 | 6.63e-08 |
| MTX1 | 600605 | 1 | 155208699 | 155213824 | -5.36 | 8.39e-08 |
| KLHDC7A | | 1 | 18480982 | 18486126 | -5.20 | 2.01e-07 |
| CASP8 | 601763 | 2 | 201233443 | 201287711 | -6.10 | 1.04e-09 |
| SLC4A7 | 603353 | 3 | 27372721 | 27484420 | -14.30 | 2.34e-46 |
| ZBTB38 | 612218 | 3 | 141324213 | 141449792 | -7.68 | 1.54e-14 |
| LINC00886 | | 3 | 156747346 | 156817062 | 5.89 | 3.96e-09 |
| CMSS1 | | 3 | 99817834 | 100181732 | 5.44 | 5.31e-08 |
| L3MBTL3 | 618844 | 6 | 130013699 | 130141451 | 6.32 | 2.67e-10 |
| PIDD1 | 605247 | 11 | 799191 | 809646 | -6.90 | 5.21e-12 |
| CFL1 | 601442 | 11 | 65823022 | 65861946 | -5.41 | 6.46e-08 |
| SPTY2D1 | | 11 | 18606401 | 18634791 | 4.80 | 1.57e-06 |
| CCDC91 | 617366 | 12 | 28133249 | 28581511 | -4.82 | 1.45e-06 |
| RCCD1 | 617997 | 15 | 90955796 | 90963125 | -5.98 | 2.18e-09 |
| PRC1-AS1 | | 15 | 90972860 | 90988624 | 5.92 | 3.19e-09 |
| CBX8 | 617354 | 17 | 79792132 | 79801683 | 5.87 | 4.35e-09 |
| ANKLE1 | 619348 | 19 | 17281645 | 17287646 | -5.26 | 1.45e-07 |
| PLA2G6 | 603604 | 22 | 38111495 | 38205690 | -5.16 | 2.46e-07 |

**Table S6: TWAS risk genes of breast cancer by PrediXcan that were identified by previous GWAS.**

| Gene | MIM | CHR | Start | End | Z-score | P-value |
|---|---|---|---|---|---|---|
| NSF | 601633 | 17 | 46590669 | 46757464 | 4.95 | 7.44e-07 |

**Table S7: TWAS risk genes of ovarian cancer by PrediXcan that were identified by previous GWAS.**

| | Traits | |
|---|---|---|
| TWAS Method | Breast Cancer | Ovarian Cancer |
| TIGAR | 88 | 37 |
| PrediXcan | 56 | 4 |

**Table S8: Number of significant TWAS risk genes identified by PrediXcan and TIGAR.**

| | Traits | |
|---|---|---|
| TWAS Method | Breast Cancer | Ovarian Cancer |
| TIGAR | 31 | 4 |
| PrediXcan | 32 | 2 |

**Table S9: Number of independently significant TWAS risk genes identified by PrediXcan and TIGAR.**

| | | | | | TIGAR | | PrediXcan | |
|---|---|---|---|---|---|---|---|---|
| Gene | MIM | CHR | Start | End | Breast | Ovarian | Breast | Ovarian |
| PRC1-AS1[a,d] | | 15 | 90972860 | 90988624 | X | X | X | X |
| LRRC37A4P[b,d] | | 17 | 45506741 | 45550335 | X | X | X | X |
| DND1P1[b,d] | | 17 | 45585871 | 45586929 | X | X | X | |
| LINC02210[b,d] | | 17 | 45620328 | 45655156 | X | X | | X |
| KLHDC7A[a] | | 1 | 18480982 | 18486126 | X | | X | |
| CH17-437K3.1[b] | | 1 | 121396754 | 121463129 | X | | X | |
| ASH1L[a,d] | 607999 | 1 | 155335268 | 155562807 | X | | X | |
| CASP8[a] | 601763 | 2 | 201233443 | 201287711 | X | | X | |
| ALS2CR12[b] | | 2 | 201288271 | 201357398 | X | | X | |
| SLC4A7[a,d] | 603353 | 3 | 27372721 | 27484420 | X | | X | |
| PSMD6-AS2[b] | | 3 | 64004022 | 64012148 | X | | X | |
| SLC22A5[b] | 603377 | 5 | 132369752 | 132395614 | X | | X | |
| PDLIM4[b] | 603422 | 5 | 132257671 | 132273454 | X | | X | |
| ANKRD55[b,d] | 615189 | 5 | 56099678 | 56233359 | X | | X | |
| C5orf56[b] | | 5 | 132410636 | 132488702 | X | | X | |
| L3MBTL3[a] | 618844 | 6 | 130013699 | 130141451 | X | | X | |
| PIDD1[a] | 605247 | 11 | 799191 | 809646 | X | | X | |
| AP006621.5[b] | | 11 | 777578 | 784297 | X | | X | |
| CCDC91[a] | 617366 | 12 | 28133249 | 28581511 | X | | X | |
| RCCD1[a,c] | 617997 | 15 | 90955796 | 90963125 | X | | X | |
| CBX8[a] | 617354 | 17 | 79792132 | 79801683 | X | | X | |
| LRRC25[b] | 607518 | 19 | 18391144 | 18397617 | X | | X | |
| UBE2MP1 | | 16 | 35169692 | 35170241 | X | X | | |
| MAPK8IP1P2[b,d] | | 17 | 45600869 | 45602340 | X | X | | |
| CRHR1[b,d] | 122561 | 17 | 45784280 | 45835828 | X | X | | |
| RP11-707O23.1[b,d] | | 17 | 45592621 | 45593369 | X | X | | |
| RP11-259G18.1[b,d] | | 17 | 46267037 | 46268694 | X | X | | |
| AC091132.1[b,d] | | 17 | 45452844 | 45464065 | X | X | | |
| ARHGAP27[a,d] | 610591 | 17 | 45393902 | 45434421 | X | X | | |
| MAPT[a,d] | 157140 | 17 | 45894382 | 46028334 | X | X | | |
| LRRC37A2[b,d] | 616556 | 17 | 46511511 | 46553449 | X | X | | |
| RP11-259G18.3[b,d] | | 17 | 46259551 | 46260606 | X | X | | |
| KANSL1-AS1[b,d] | | 17 | 46193576 | 46196723 | X | X | | |
| FAM215B[b,d] | | 17 | 46558830 | 46562795 | X | X | | |
| FRG1EP | | 20 | 29480147 | 29497179 | X | X | | |
| NSF[b,c] | 601633 | 17 | 46590669 | 46757464 | | X | | X |

*a* known GWAS risk gene of breast cancer.

*b* within 1MB of a known GWAS risk gene of breast cancer.

*c* known GWAS risk gene of ovarian cancer.

*d* within 1MB of a known GWAS risk gene of ovarian cancer.

**Table S10: Significant TWAS risk genes identified by PrediXcan and TIGAR for breast and ovarian cancer.** The "X" symbol denotes a significant TWAS risk gene for the corresponding trait by the corresponding method.

| Method | Traits | Gene | CHR | Start | End | Z-score | P-value |
|---|---|---|---|---|---|---|---|
| TIGAR | Breast | KLHL25 | 15 | 85759323 | 85795030 | -4.73 | 2.22e-06 |
| TIGAR | Breast | UBE2MP1 | 16 | 35169692 | 35170241 | -5.31 | 1.13e-07 |
| TIGAR | Ovarian | UBE2MP1 | 16 | 35169692 | 35170241 | 5.77 | 7.88e-09 |
| TIGAR | Breast | ANKRD20A21P | 20 | 30656033 | 30723932 | -5.22 | 1.83e-07 |
| TIGAR | Breast | FRG1EP | 20 | 29480147 | 29497179 | 5.39 | 6.95e-08 |
| TIGAR | Ovarian | FRG1EP | 20 | 29480147 | 29497179 | -4.99 | 6.19e-07 |
| PrediXcan | Breast | RP11-337N6.3 | 2 | 177317715 | 177318471 | 5.46 | 4.82e-08 |
| PrediXcan | Breast | EFCC1 | 3 | 129001629 | 129040742 | 4.84 | 1.32e-06 |

$*$ No MIM identifier available for any gene in this table.

**Table S11: Significant TWAS risk genes of breast and ovarian cancer identified by PrediXcan and TIGAR but not by previous GWAS.**

### Supplemental Text

#### Text S1 Nonparametric Bayesian DPR model

Given reference genotype matrix $\mathbf{G}$ of SNPs within ±1MB of the transcription start sites of the target gene $g$ and expression quantitative trait $\mathbf{E}_g$ of the target gene, both centered at 0, the nonparametric Bayesian Dirichlet Process Regression (DPR) model assumes

$$\mathbf{E}_g = \mathbf{G}\mathbf{w}_g + \boldsymbol{\varepsilon}_g, \quad \boldsymbol{\varepsilon}_g \sim \mathrm{N}(0, \sigma_\varepsilon^2 \mathbf{I}), \quad \sigma_\varepsilon^2 \sim \mathrm{IG}(a_\varepsilon, b_\varepsilon), \tag{S1}$$

where $\mathbf{w}$ denotes the *cis*-eQTL effect size vector. Note that such gene expression prediction model need to be fitted per gene using the SNP genotype data ±1MB of the test gene as predictors.

The effect-size of each *cis*-eQTL ($w_\mathrm{i}$; $i = 1, \ldots, p$) is assumed to follow a normal prior $\mathrm{N}(0, \sigma_w^2)$ with an effect-size variance $\sigma_w^2$ that is assumed to follow a Dirichlet Process (DP) prior $D$ with an inverse gamma (IG) base distribution and a concentration parameter $\xi$.

$$w_\mathrm{i} \sim \mathrm{N}(0, \sigma_w^2), \quad \sigma_w^2 \sim D, \quad D \sim \mathrm{DP}(\mathrm{IG}(a, b), \xi).$$

Effect-size variance $\sigma_w^2$ can be viewed as a latent variable that can be integrated out in order to derive the following equivalent DP normal mixture model:

$$w_\mathrm{i} \sim \sum_{k=0}^{+\infty} \pi_k \mathrm{N}(0, \sigma_k^2), \quad \sigma_k^2 \sim \mathrm{IG}(a_k, b_k), \quad \pi_k = v_k \prod_{l=0}^{k-1} (1 - v_l), \quad v_k \sim \mathrm{Beta}(1, \xi).$$

The resulting mixture normal prior is the weighted sum of an infinite number of normal distributions $\mathrm{N}(0, \sigma_k^2)$ with weights $\pi_k$ determined by $v_l$, $l = (0, \cdots, k-1, k)$ with a Beta prior. The number of components with non-zero weights in the mixture normal prior is determined by the concentration parameter $\xi$, which is assumed to follow a $\mathrm{Gamma}(a_\xi, b_\xi)$ hyper prior.

Non-informative priors for ($\sigma_k^2$, $\sigma_\epsilon^2$, $\xi$) are induced by setting hyper parameters $a_k$, $b_k$, $a_\epsilon$, $b_\epsilon$, $b_\xi = 0.1$ and $a_\xi = 1$. That is, the parameters ($\sigma_k^2$, $\sigma_\epsilon^2$, $\xi$) will be adaptively estimated from the data and the nonparametric prior on $w_i$ will be data-driven.

The variational Bayesian algorithm [1, 2], an approximation for the Gibbs Sampling MCMC algorithm [3] with greater computational efficiency, is then used to adaptively estimate the parameters ($\sigma_k^2$, $\sigma_\epsilon^2$, $\xi$) and obtain the Bayesian posterior estimates for $\mathbf{w}$. More details about the Bayesian inference algorithm can be found in the Supplemental Note of the TIGAR paper [4].

#### Text S2 TWAS with GWAS data

With both individual-level and summary-level GWAS data, TIGAR-V2 implements both Burden [5, 6] and Variance-Component [7] test for TWAS, where eQTL effect size estimates $\widehat{\mathbf{w}}$ from the gene expression prediction model of the test gene are taken as variant weights. Note that here the eQTL effect size estimates $\widehat{\mathbf{w}}$ are specific to the test gene and specific to the tissue type of the reference transcriptomic data.

#### Text S2.1 TWAS with individual-level GWAS data

With the Bayesian posterior estimates of the eQTL effect sizes $\widehat{\mathbf{w}}$ and individual-level GWAS data of a test cohort, the genetically regulated gene expression (GReX) of the test gene with genotype matrix $\mathbf{G}_{\text{test}}$ is then imputed by

$$\widehat{\mathbf{GReX}}_g = \mathbf{G}_{\text{test}} \widehat{\mathbf{w}}.$$

The Burden type gene-based TWAS [5, 6] would basically test the association between the imputed $\widehat{\mathbf{GReX}}_g$ values and the phenotype of interest based on a general linear regression model. The Variance-Component TWAS [7] would basically use the sequence kernel association test (SKAT) framework [8] with variant weights provided by Bayesian eQTL weights $\widehat{\mathbf{w}}$.

#### Text S2.2 TWAS with summary-level GWAS data

##### Burden TWAS

In the following derivations, we show that the FUSION Z-score statistic [9] as given by Equation (S2) will lead to inflated false positive findings and the S-PrediXcan test statistic [10] as given by Equation (S3) should be used, if $\widehat{\mathbf{w}}$ are estimated using non-standardized reference data (i.e., centered gene expression and genotype data as described in Text S1). We also show that both FUSION and S-PrediXcan test statistics are equivalent if $\widehat{\mathbf{w}}$ are estimated using standardized reference data.

$$\widetilde{Z}_{g,\text{FUSION}} = \frac{\sum_{l=1}^{m} (\widehat{w_l} Z_l)}{\sqrt{\widehat{\mathbf{w}}' \mathbf{V} \widehat{\mathbf{w}}}}, \quad \mathbf{V} = \text{Corr}(\mathbf{G}_0) \tag{S2}$$

$$\widetilde{Z}_{g,\text{S-PrediXcan}} = \frac{\sum_{l=1}^{m} (\widehat{w_l} \widehat{\sigma_l} Z_l)}{\sqrt{\widehat{\mathbf{w}}' \mathbf{V} \widehat{\mathbf{w}}}}, \quad \widehat{\sigma_l^2} = \text{Var}(\mathbf{G_{0l}}), \quad \mathbf{V} = \text{Cov}(\mathbf{G}_0). \tag{S3}$$

Here, $Z_l$ denotes the Z-score statistic values from single variant GWAS tests (i.e., summary-level GWAS data). The required linkage disequilibrium (LD) covariance matrix (or correlation matrix for FUSION test statistic) among test cis-SNPs ($\mathbf{V}$), and the variance of test cis-SNPs ($\widehat{\sigma_l^2} = \text{Var}(\mathbf{G_{0l}})$) can be obtained from reference genotype data ($\mathbf{G}_0$) such as 1000 genome [11] and GTEx V8 [12].

##### Test Z-score statistic for Burden TWAS

Consider the above assumed gene expression prediction model as in Equation S1, and the following model of phenotype assumed by Burden TWAS

$$\mathbf{Y} = \gamma \mathbf{E}_{\text{test}} + \epsilon_{\mathbf{Y}}, \ \epsilon_{\mathbf{Y}} \sim N(0, \sigma_{\epsilon,Y}^2 \mathbf{I}), \tag{S4}$$

where $\mathbf{Y}$ denotes the phenotype vector and $\mathbf{E}_{\text{test}}$ denotes the GReX values of the test gene. In Burden TWAS with individual-level GWAS data of phenotype vector $\mathbf{Y}$ and test genotype data $\mathbf{G}_{\text{test}}$, it is basically testing $H_0 : \gamma = 0$ by replacing $\mathbf{E}_{\text{test}}$ in Equation S4 by $\widehat{\mathbf{GReX}} = \mathbf{G}_{\text{test}} \widehat{\mathbf{w}}$. That is,

$$\mathbf{Y} = \gamma(\mathbf{G}_{\text{test}} \widehat{\mathbf{w}}) + \epsilon_{\mathbf{Y}} = \mathbf{G}_{\text{test}}(\gamma \widehat{\mathbf{w}}) + \epsilon_{\mathbf{Y}} = \mathbf{G}_{\text{test}} \beta + \epsilon_{\mathbf{Y}},$$

where the SNP effect sizes on phenotype are assumed to be a linear function of the eQTL weights $\widehat{\mathbf{w}}$, $\beta = \gamma \widehat{\mathbf{w}}$. Generally, summary-level GWAS data provide marginal lease square estimates for $\beta$,

$$\widehat{\beta_l} = (\mathbf{G}'_{\text{test},l} \mathbf{G}_{\text{test},l})^{-1} (\mathbf{G}'_{\text{test},l} \mathbf{Y}), \ \text{Var}(\widehat{\beta_l}) = \sigma_Y^2 (\mathbf{G}'_{\text{test},l} \mathbf{G}_{\text{test},l})^{-1}, \ l = (1, \cdots, m), \tag{S5}$$

where $\sigma_Y^2 = \text{Var}(\mathbf{Y})$ denotes the variance of phenotype.

Here, we assume all reference data $\mathbf{E_g}, \mathbf{G}$ and test data $\mathbf{Y}, \mathbf{E}_{\text{test}}, \mathbf{G}_{\text{test}}$ are centered to have zero column means. The least square estimate of $\gamma$ in Equation S4 is given by

$$\widehat{\gamma} = (\mathbf{E}'_{\text{test}}\mathbf{E}_{\text{test}})^{-1}(\mathbf{E}'_{\text{test}}\mathbf{Y}), \ \text{Var}(\widehat{\gamma}) = \sigma_Y^2(\mathbf{E}'_{\text{test}}\mathbf{E}_{\text{test}})^{-1}.$$

Then the Burden TWAS Wald Z-score statistic for testing $H_0 : \gamma = 0$ is given by

$$\begin{aligned} Z_{score,\gamma} &= \frac{\widehat{\gamma}}{\sqrt{\text{Var}(\widehat{\gamma})}} = \frac{(\mathbf{E}'_{\text{test}}\mathbf{E}_{\text{test}})^{-1}(\mathbf{E}'_{\text{test}}\mathbf{Y})}{\sqrt{\sigma_Y^2(\mathbf{E}'_{\text{test}}\mathbf{E}_{\text{test}})^{-1}}} = \frac{\mathbf{E}'_{\text{test}}\mathbf{Y}}{\sqrt{\sigma_Y^2}\sqrt{\mathbf{E}'_{\text{test}}\mathbf{E}_{\text{test}}}} \\ &= \frac{\widehat{\mathbf{w}}'(\mathbf{G}'_{\text{test}}\mathbf{Y})}{\sqrt{\sigma_Y^2}\sqrt{\widehat{\mathbf{w}}'(\mathbf{G}'_{\text{test}}\mathbf{G}_{\text{test}})\widehat{\mathbf{w}}}} = \frac{\sum_{l=1}^m \left(\widehat{w}_l(\mathbf{G}'_{\text{test},l}\mathbf{Y})\right)}{\sqrt{\sigma_Y^2}\sqrt{\widehat{\mathbf{w}}'(\mathbf{G}'_{\text{test}}\mathbf{G}_{\text{test}})\widehat{\mathbf{w}}}}, \end{aligned} \tag{S6}$$

where $\mathbf{E}_{\text{test}} = \widehat{\mathbf{w}}\mathbf{G}'_{\text{test}}$.

From the marginal least square estimates of SNP effect sizes on phenotype as in Equation S5, we can derive the following estimates of the required quantities in the Wald's Z-score statistic of $\gamma$,

$$\mathbf{G}'_{\text{test},l}\mathbf{Y} = (\mathbf{G}'_{\text{test},l}\mathbf{G}_{\text{test},l})\widehat{\beta}_l; \ \mathbf{G}'_{\text{test}}\mathbf{G}_{\text{test}} = (n-1)\mathbf{V}; \ \mathbf{G}'_{\text{test},l}\mathbf{G}_{\text{test},\mathbf{l}} = (n-1)\mathbf{V}_{(\mathbf{l},\mathbf{l})},$$

where genotype covariance matrix $\mathbf{V}$ can be approximated by $\text{Cov}(\mathbf{G_0})$ with genotype data $\mathbf{G_0}$ from a reference panel of the same ancestry. That is,

$$\begin{aligned} Z_{\text{score},\gamma} &= \frac{\sum_{l=1}^m \left(\widehat{w}_l(\mathbf{G}'_{\text{test},l}\mathbf{G}_{\text{test},l})\widehat{\beta}_l\right)}{\sqrt{\sigma_Y^2}\sqrt{(n-1)(\widehat{\mathbf{w}}'\mathbf{V}\widehat{\mathbf{w}})}} = \frac{(n-1)\sum_{l=1}^m \left(\widehat{w_l}\mathbf{V}_{(\mathbf{l},\mathbf{l})}\widehat{\beta}_l\right)}{\sqrt{\sigma_Y^2}\sqrt{(n-1)}\sqrt{\widehat{\mathbf{w}}'\mathbf{V}\widehat{\mathbf{w}}}} \\ &= \frac{\sqrt{(n-1)}\sum_{l=1}^m \left(\widehat{w}_l\mathbf{V}_{(\mathbf{l},\mathbf{l})}\widehat{\beta}_l\right)}{\sqrt{\sigma_Y^2}\sqrt{\widehat{\mathbf{w}}'\mathbf{V}\widehat{\mathbf{w}}}}, \end{aligned} \tag{S7}$$

where $n$ denotes the sample size of the test cohort.

#### FUSION Z-Score Statistic

Note that if the GWAS summary statistics were derived from standardized genotype matrix $\mathbf{G}_{\text{test}}$ and phenotype vector $\mathbf{Y}$, then $\mathbf{V}$ denotes the genotype correlation matrix, $\mathbf{V}_{(\mathbf{l},\mathbf{l})} = 1$, $\sigma_Y^2 = 1$, and the GWAS Z-score statistic for SNP $l$ can be denoted as

$$Z_l = \frac{\widehat{\beta}_l}{\sqrt{\text{Var}(\widehat{\beta}_l)}} = \frac{\widehat{\beta}_l}{1/\sqrt{(n-1)}} = \sqrt{(n-1)}\widehat{\beta}_l.$$

Thus,

$$Z_{\text{score},\gamma} = \frac{\sum_{l=1}^m (\widehat{w}_l Z_l)}{\sqrt{\widehat{\mathbf{w}}'\mathbf{V}\widehat{\mathbf{w}}}},$$

which is the FUSION test Z-score statistic as in Equation S2. Note that the SNP effect sizes here are assumed to be a linear function of the eQTL effect sizes estimated from the gene expression prediction model as in Equation S1, $\beta = \gamma\widehat{\mathbf{w}}$. That is, to make FUSION test Z-statistic a valid test for Burden TWAS, the eQTL effect size estimates $\widehat{\mathbf{w}}$ is also assumed to be derived from standardized gene expression data $\mathbf{E_g}$ and reference genotype data $\mathbf{G}$.

#### S-PrediXcan Z-Score statistic

In our studies, we noticed inflation of false positive findings by Burden TWAS if FUSION test Z-score statistic is used along with $\widehat{\mathbf{w}}$ estimated from non-standardized reference gene expression and genotype data. As assumed in Bayesian DPR model implemented in the TIGAR method, reference gene expression and genotype data are only centered to have zero column means but not standardized to have unit variance. In this scenario, the Burden TWAS assumes the SNP effect sizes on phenotype ($\beta = \gamma \widehat{\mathbf{w}}$ ) are not derived with standardized phenotype and test genotype data. Then the single variant Z-score statistics in GWAS data are given by

$$Z_l = \frac{\widehat{\beta_l}}{\sqrt{\mathrm{Var}(\widehat{\beta_l})}} = \frac{\widehat{\beta_l}}{\sqrt{\sigma_Y^2 (\mathbf{G}'_{\mathrm{test},l} \mathbf{G}_{\mathrm{test},l})^{-1}}} = \frac{\sqrt{(n-1)\mathbf{V}_{(\mathbf{1},\mathbf{1})}}\widehat{\beta_l}}{\sqrt{\sigma_Y^2}}.$$

Then the Burden TWAS Z-score statistic in Equation S7 is equivalent to the S-PrediXcan test statistic [10] as given by Equation (S3),

$$\begin{aligned} Z_{\mathrm{score},\gamma} &= \frac{\sqrt{(n-1)}\sum_{l=1}^m \left(\widehat{w}_l \mathbf{V}_{(\mathbf{1},\mathbf{1})} \widehat{\beta_l}\right)}{\sqrt{\sigma_Y^2}\sqrt{\widehat{\mathbf{w}}'\mathbf{V}\widehat{\mathbf{w}}}} = \frac{\sum_{l=1}^m \left(\widehat{w}_l \sqrt{\mathbf{V}_{(\mathbf{1},\mathbf{1})}} (\sqrt{(n-1)}\sqrt{\mathbf{V}_{(\mathbf{1},\mathbf{1})}}\widehat{\beta_l})\right)}{\sqrt{\sigma_Y^2}\sqrt{\widehat{\mathbf{w}}'\mathbf{V}\widehat{\mathbf{w}}}} \\ &= \frac{\sum_{l=1}^m \left(\widehat{w}_l \sqrt{\mathbf{V}_{(\mathbf{1},\mathbf{1})}} Z_l\right)}{\sqrt{\widehat{\mathbf{w}}'\mathbf{V}\widehat{\mathbf{w}}}} \approx \frac{\sum_{l=1}^m \left(\widehat{w}_l \widehat{\sigma_l^2} Z_l\right)}{\sqrt{\widehat{\mathbf{w}}'\mathbf{V}\widehat{\mathbf{w}}}}, \end{aligned} \tag{S8}$$

where $\widehat{\sigma_l^2} \approx \mathrm{Var}(\mathbf{G_{0,l}})$ denotes the genotype variance of SNP $l$.

#### Variance-Component TWAS test statistic

With summary-level GWAS data, Variance-Component TWAS test can still be conducted. Details about the test statistic derivations are provided in [7]. Variance-Component TWAS test is recommended if the assumption of the linear relationship between the SNP effect sizes on phenotype and eQTL weights ($\beta = \gamma \widehat{\mathbf{w}}$) is violated.
